# Stomatal regulators are co-opted for seta development in the astomatous liverwort *Marchantia polymorpha*

**DOI:** 10.1101/2022.04.29.489008

**Authors:** Kenta C. Moriya, Makoto Shirakawa, Jeanne Loue-Manifel, Yoriko Matsuda, Yen-Ting Lu, Kentaro Tamura, Yoshito Oka, Tomonao Matsushita, Ikuko Hara-Nishimura, Gwyneth Ingram, Ryuichi Nishihama, Justin Goodrich, Takayuki Kohchi, Tomoo Shimada

## Abstract

The evolution of special types of cells requires the acquisition of new gene regulatory networks controlled by transcription factors (TFs). In stomatous plants, a TF module formed by subfamilies Ia and IIIb basic helix-loop-helix TFs (Ia-IIIb bHLH) regulates stomatal formation; however, how this module evolved during land plant diversification remains unclear. Here, we show that, in the astomatous liverwort *Marchantia polymorpha*, a Ia-IIIb bHLH module regulates the development of a unique sporophyte tissue, the seta, which is found in mosses and liverworts. The sole Ia bHLH gene, Mp*SETA*, and a IIIb bHLH gene, Mp*ICE2*, regulate the cell division and/or differentiation of seta lineage cells. MpSETA can partially replace the stomatal function of Ia bHLH TFs in *Arabidopsis thaliana*, suggesting that a common regulatory mechanism underlies the setal and stomatal formation. Our findings reveal the co-option of a Ia-IIIb bHLH TF module for regulating cell fate determination and/or cell division of distinct types of cells during land plant evolution.

---

Land plants developed unique types of cells and tissues to adapt to the terrestrial environment during evolution^1,2^. The acquisition of novel cells or tissues leading to complex body plans is related to the diversification of transcription factors (TFs)^3,4^. Basic helix-loop-helix (bHLH) TFs represent a TF superfamily that plays key roles in cell fate determination and cell division during eukaryote development. In land plants, the number of genes encoding bHLH TFs has increased compared with those of chlorophyte and charophyte algae, suggesting that bHLH TFs are involved in terrestrial adaptation^5^. For example, the acquisition of stomata, a special tissue for gas exchange on the epidermis, is an important adaptation of plants to terrestrial environments; previous studies have revealed that stomatal formation is regulated by bHLH TFs as master TFs^6–9^. In *Arabidopsis thaliana*, three TFs belonging to subfamily Ia (SPEECHLESS [SPCH], MUTE, and FAMA)^6–8^ form heterodimers with subfamily IIIb TFs (ICE1, also known as SCREAM [SCRM], and ICE2, also known as SCRM2) to promote the stomatal formation by regulating downstream gene expression^9^. The molecular mechanism of stomatal formation by Ia and IIIb bHLH TFs is conserved in the moss *Physcomitrium patens*, in which stomata play an important role in spore dispersal by promoting dehydration and dehiscence of the sporangium^10,11^.

Recent studies have revealed that a gene encoding a Ia bHLH is present in the genome of the astomatous liverwort *Marchantia polymorpha*^4,12–14^. Despite the importance of Ia bHLH in stomatal development, its function in plants without stomata remains unexplored. Here, we report that the Ia bHLH protein, designated as MpSETA, is a master regulator of the formation of the seta, which is a diploid tissue involved in long-distance spore dispersal in *M. polymorpha*. Furthermore, we show that Ia and IIIb bHLH positively regulate setal formation by heterodimerization, similar to their role in stomatal formation in other land plants. This outcome advances our understanding of the mechanisms of the evolution of plant tissue formation while providing new insights into the co-option of gene expression regulatory networks (GRNs).

## Results

### MpSETA is the sole Ia bHLH protein in *M. polymorpha*

To identify Ia bHLH coding genes in *M. polymorpha*, we constructed a phylogenetic tree of Ia bHLH proteins from various plant species using a bHLH domain and C-terminal conserved domain called SMF (also known as the ACT-like domain)^15,16^ (Fig. 1a and Extended Data Fig. 1a). Our phylogenetic analysis suggested that MpBHLH35 (Mp2g04160) is the sole Ia bHLH in *M. polymorpha* (Fig. 1a). We named this gene Mp*SETA* based on its specific expression in the seta tissue of the sporophyte (see below). Multiple alignments of the bHLH proteins revealed that the amino acid residues predicted to be important for E-box (CANNTG) binding and bHLH dimerization are highly conserved in MpSETA, although the amino acid sequence of the bHLH domain of MpSETA is relatively divergent compared to other Ia bHLH (Extended Data Fig. 1b). Additionally, we found partial sequences of two Mp*SETA* related genes (Lc*SETA1* and Lc*SETA2*) in the genome of the Marchantiidae liverwort *Lunularia cruciata*^17^, although there is no evidence that these putative MpSETA-like genes are expressed and are functional in this species (Fig.1a and Extended Data Fig. 1b-c). The amino acid sequences of the bHLH and SMF domains are well conserved between MpSETA and LcSETA1. Thus, we can conclude that Ia bHLH genes are conserved in the genome of Marchantiidae liverworts.

**Fig. 1.**
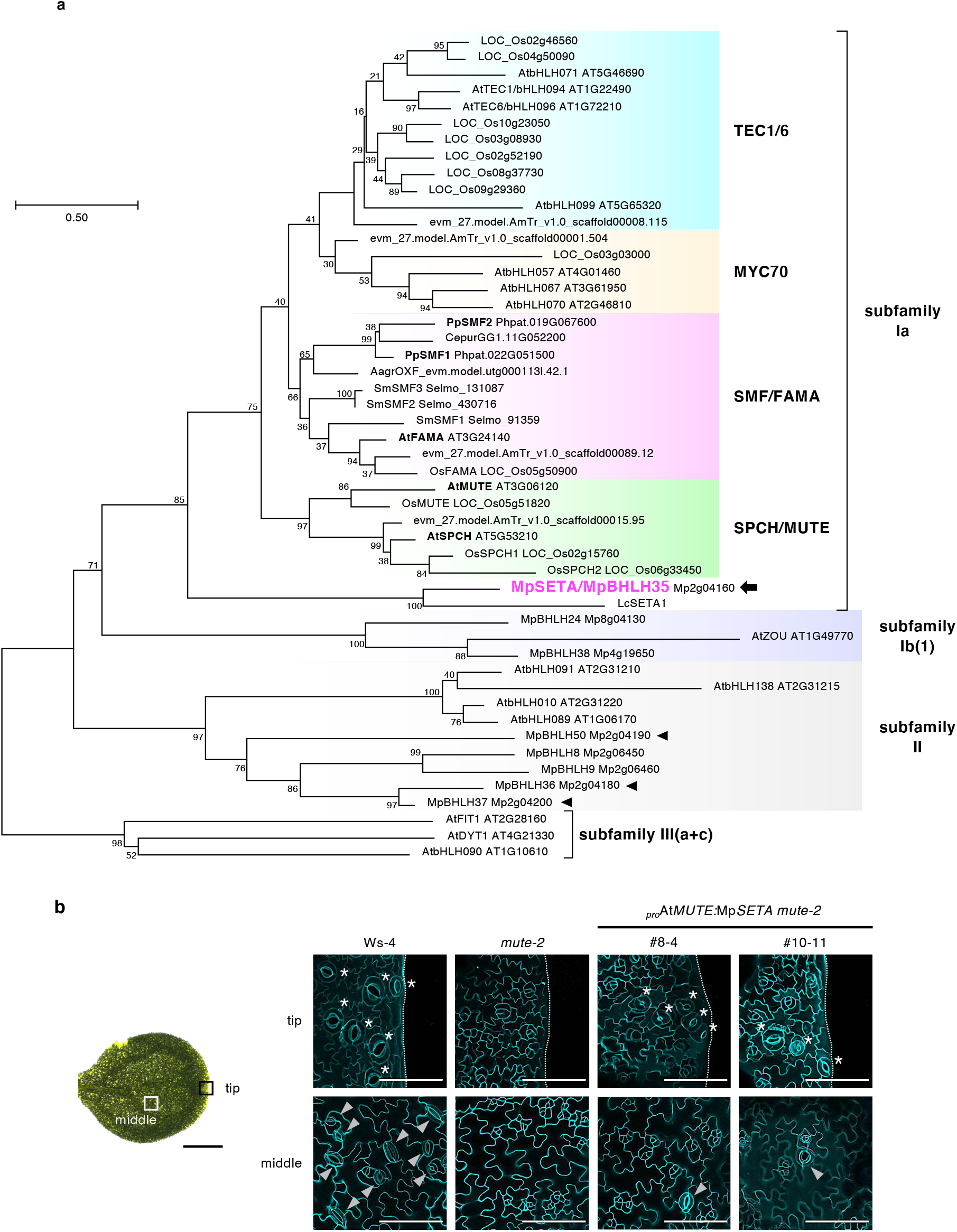
MpSETA is the only bHLH transcription factor that belongs to subfamily Ia in *M. polymorpha*. **a**, A maximum-likelihood bHLH phylogenetic tree of subfamilies Ia, Ib(1) (purple), II (gray), and III(a+c) (outgroup) is shown. Numbers at branches indicate bootstrap values calculated from 1,000 replicates. Ia bHLHs are divided into four groups: TEC1/6 clade (light blue), MYC70 clade (cream-yellow), SMF/FAMA clade (magenta), and SPCH/MUTE clade (green). Species are abbreviated as follows: Mp, *M. polymorpha* (liverwort); Lc, *L. cruciata* (liverwort); Pp, *P. patens* (moss); Cepur, *Ceratodon purpureus* (moss); Aagr, *Anthoceros agrestis* (hornwort); Sm, *Selaginella moellendorffii* (lycophyte); AmTr, *Amborella trichopoda* (basal angiosperm); Os, *Oryza sativa* (monocot); At, *A. thaliana* (dicot). An arrow indicates MpSETA/MpBHLH35 (Mp2g04160), and arrowheads indicate bHLH TFs mentioned as FAMA-like bHLH TFs in a previous study^14^. Amino acid sequences from only *A. thaliana* and *M. polymorpha* were used for subfamilies Ib(1), II, and III(a+c). **b**, Confocal images of *A. thaliana* abaxial cotyledons of wild type (Ws-4), *mute-2*, and _*pro*_At*MUTE:*Mp*SETA mute-2* at 9 days after stratification (DAS). The upper and lower panels show the middle and tip areas of the cotyledons, respectively (left image). Arrowheads and asterisks indicate stomata and hydathode pores, respectively. Bars, 100 μm (**b**, confocal images), and 1 mm (**b**, left).

Because the amino acid sequence of MpSETA is divergent, whether MpSETA shares similar properties with the other Ia bHLH proteins is unclear. Therefore, we investigated whether MpSETA can act as a stomatal regulator replacing AtSPCH, AtMUTE, or AtFAMA. In this context, Mp*SETA* was expressed under the native promoters of At*SPCH*, At*MUTE*, and At*FAMA* in *spch-3, mute-2*, and *fama-1* backgrounds, respectively (Fig. 1b and Extended Data Fig. 2a,b). Even though *mute-2* results in arrested meristemoids (self-renewing stomatal precursors), a few stomata were formed in *mute-2* expressing Mp*SETA* (Fig. 1b) (2.67 ± 1.56 and 2.33 ± 1.72 per abaxial side of the cotyledon in lines #8-4 and #10-11, respectively [mean ± s.d.; *n* = 12]). Notably, hydathode pores (a modified form of stomatal pores) were often found in these lines. This might be due to the high expression activity of the At*MUTE* promoter in the hydathode of cotyledons^18^. Mp*SETA* also exhibited the potential to rescue *fama-1. A. thaliana fama* mutant displays caterpillar-like stomatal-lineage cells called “*fama* tumors,” where terminal symmetric division occurs more than once^8^. In *fama-1* expressing Mp*SETA*, excess cell divisions in the stomatal lineage were suppressed, although no mature stomata were formed (Extended Data Fig. 2b,c). Neither stomata nor stomatal-lineage cells were found in *spch-3* expressing Mp*SETA* (Extended Data Fig. 2a). These results, showing that Mp*SETA* is partially functional in stomatal cell division and differentiation in *A. thaliana*, suggest that it can interact with AtICE1 and AtSCRM2. Therefore, we tested this ability in yeast two-hybrid assays and bimolecular fluorescent complementation (BiFC) assays. We found that MpSETA physically interacted with AtICE1 and AtSCRM2 (Extended Data Fig. 2d,e). Thus, our findings indicate that MpSETA from the astomatous liverwort is a bona fide Ia bHLH TF, although its amino acid sequence is divergent.

### Mp*SETA* is preferentially expressed in developing sporophyte

To investigate the expression pattern of Mp*SETA* in *M. polymorpha*, we reanalyzed the public RNA-seq dataset from several organs^4,19–22^ and found that Mp*SETA* was preferentially expressed in the diploid sporophyte, whereas its expression level was low in the haploid gametophyte (Fig. 2a).

**Fig. 2.**
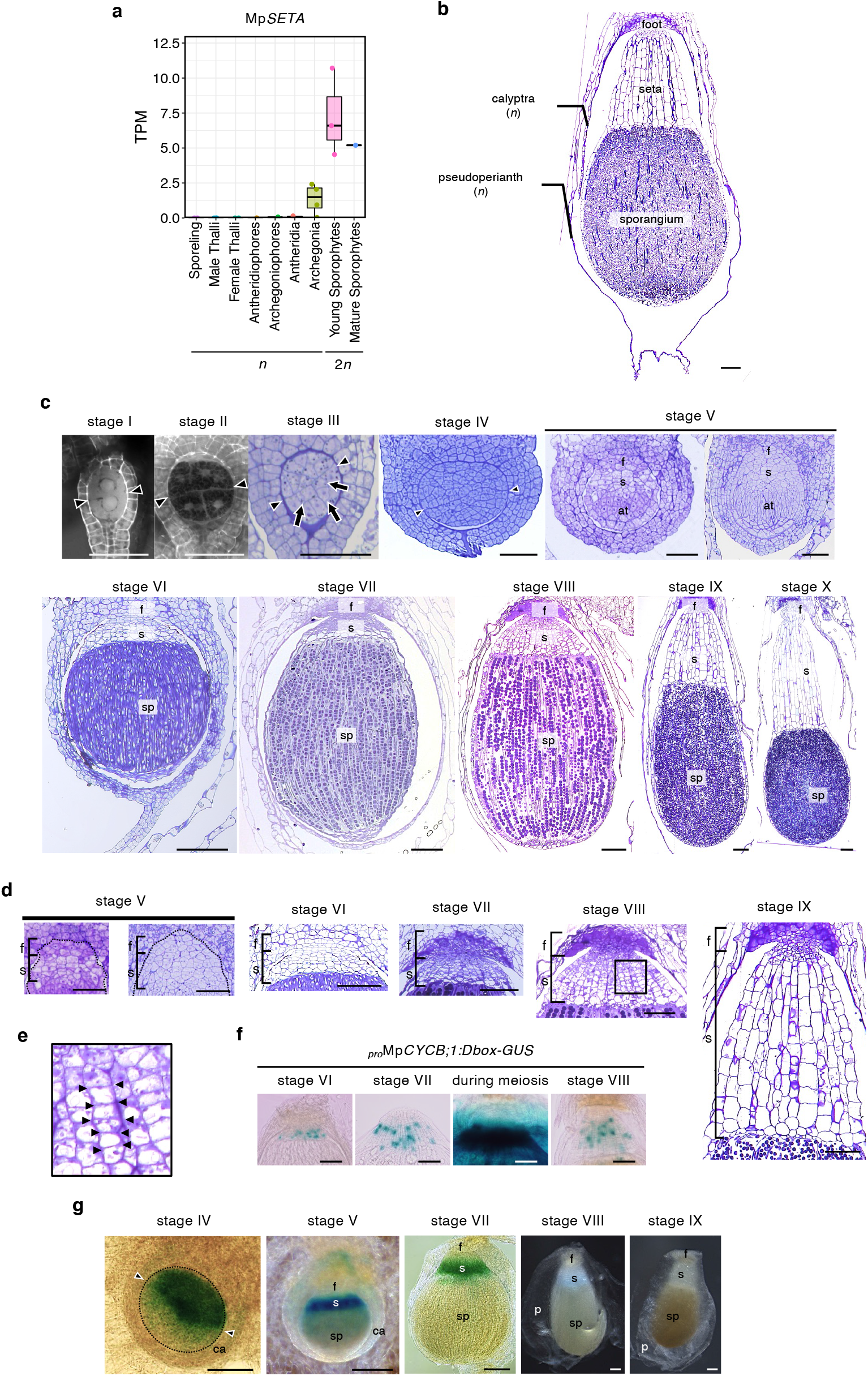
Mp*SETA* is preferentially expressed in developing sporophyte. **a**, Box plot showing the expression profiles of Mp*SETA* across nine *M. polymorpha* tissues. The Y-axis shows transcripts per million (TPM). Sporeling and thalli are vegetative gametophytic (*n*) organs. Antheridiophores and archegoniophores are haploid male and female reproductive receptacles, respectively. Antheridia and archegonia are the organs that produce sperm and egg cells, respectively. Sporophytes are diploid (2*n*) organs that develop after fertilization. **b**, Tissue section of mature *M. polymorpha* sporophyte (stage IX). **c**, Developmental stages of *M. polymorpha* sporophytes after 4–40 days’ postfertilization. (I) sporophyte differentiating an epibasal cell and a hypobasal cell, (II) 4- or 8-cell sporophyte, (III) sporophyte differentiating amphithecium and endothecium, (IV) later globular sporophyte, (V) sporophyte differentiating foot, seta, and archesporial tissues, (VI) sporophyte differentiating sporogenous cells and elaterocytes (or elater mother cell), (VII) sporophyte differentiating sporocytes (or spore mother cells), (VIII) sporophyte differentiating spore tetrads after meiosis, (IX) mature sporophyte before seta elongation, (X) mature sporophyte after seta elongation. Arrows indicate the endothecium. Arrowheads indicate the cell wall of the first cell division. **d**, Magnified images showing the foot and seta for each stage of wild-type sporophytes. The same images as used in (**c**). **e**, The formation of the cell files of seta in the wild type stage VIII sporophyte. Enlarged images of the square areas are shown in (**d**). Arrowheads indicate the plane of proliferative cell divisions. **f**, Histochemical detection of *β*-glucuronidase (GUS) activity in the sporophytes crossed with male wild type and female _*pro*_Mp*CYCB;1:Dbox-GUS*. **g**, Histochemical detection of *β*-glucuronidase (GUS) activity driven by the Mp*SETA* promoter in the developing seta region from stage IV to stage IX. f, foot; s, seta; at, archesporial tissue; sp, sporangium; ca, calyptra; p, pseudoperianth. Arrowheads indicate the cell wall of the first cell division. Bars, 50 μm (**c**, top), and 100 μm (**b, c**, bottom, **d, f** and **g**).

We examined in detail the expression pattern of Mp*SETA* by characterizing the different stages of development in the *M. polymorpha* sporophyte (Fig. 2b-c); in the wild type, sporophytes are divided into three tissues: foot, seta, and sporangium (Fig. 2b). The foot plays a key role in nutrient transport between the gametophyte and sporophyte, while the seta, which comprises files of elongated cells that form a stalk suspending the sporangium, plays a pivotal role in spore dispersal^23–25^. After late spore maturation in sporophyte development, the seta extends through the elongation of its cells and thrusts the sporangium outwards, causing it to break through the surrounding calyptra and pseudoperianth, which are tissues derived from the archegonium and archegoniophore, respectively, to protect the sporophyte during development^23–25^. As a result, spores are dispersed after sporangium dehiscence by desiccation. Based on the unique developmental events as shown in the previous studies^22,23,26^, we divided the development of the sporophyte into 10 stages, as follows (Fig. 2c,d): (I) 2-cell stage with the epibasal cell (the upper cell that forms the foot and seta) and the hypobasal cell (the lower cell that forms the sporangium); (II) 4- or 8-cell stage; (III) early-globular stage with a distinct amphithecium (the outermost tissue that forms the capsule wall) and the endothecium (the inner archesporial tissue); (IV) late-globular stage; (V) archesporial-tissue stage with the visible differentiated foot, seta, and archesporial tissues; (VI) sporogenous-cell stage with elaterocytes and sporogenous cells, which are the precursors of elaters and sporocytes (spore mother cells); (VII) sporocyte stage; (VIII) spore-tetrad stage (after meiosis); (IX) maturated stage; (X) seta-elongated stage. Note that the seta and foot are established between stages V and IX. In stage VIII, we anatomically observed proliferative (symmetric) cell divisions in the seta region (Fig. 2e). Moreover, cell division activity was detected not in the foot but rather in the seta of sporophytes between stages VII and VIII using a G2-M phase reporter line, _*pro*_Mp*CYCB;1:Dbox-GUS*^27^ (Fig. 2f). Thus, we concluded that the cell files in the seta are established by a few proliferative cell divisions of the putative “seta mother cell” in the later developmental stage.

We detected the promoter activity of Mp*SETA* in the sporophyte by generating transformants expressing the *β*-glucuronidase gene (*GUS*) under the control of the Mp*SETA* promoter (_*pro*_Mp*SETA:GUS*). GUS activity was found in developing sporophytes between stages IV and VII, especially in seta, whereas no GUS activity was found in stages VIII and IX (Fig. 2g). In gametophytic tissues, GUS signals were detected only in young antheridia (Extended Data Fig. 3). These findings suggest that Mp*SETA* is expressed early in seta development and may regulate seta cell division and/or differentiation rather than setal cell elongation.

### Mp*seta*^*ko*^ mutants show defects in setal formation in the sporophyte

We generated loss-of-function mutants of Mp*SETA* by homologous recombination-mediated gene targeting to reveal the function of Mp*SETA in vivo*^28^. Therefore, we obtained two independent Mp*SETA* knock-out lines (Mp*seta-1*^*ko*^ and Mp*seta-2*^*ko*^) in which the genomic regions encoding the bHLH domain were replaced with a hygromycin-resistance gene cassette (Extended Data Fig. 4a-c). We confirmed the loss of the full-length transcripts of the Mp*SETA* gene by reverse-transcription polymerase chain reaction (RT-PCR) analysis of homozygous mutant sporophytes produced from crosses between Mp*seta-1*^*ko*^ or Mp*seta-2*^*ko*^ males and females (Extended Data Fig. 4d). Although Mp*SETA* reporter gene expression was found in the developing antheridia (Extended Data Fig. 3), no obvious phenotype was observed during sperm formation in Mp*seta*^*ko*^ lines (Extended Data Fig. 4e), and mutant males were fertile.

We crossed males and females of Mp*seta*^*ko*^ and compared the resulting sporophytes with those of the wild type to investigate the phenotypes of Mp*seta*^*ko*^ mutants in the diploid generation. In longitudinal sections of the mature sporophytes of Mp*seta*^*ko*^, we found anatomical defects in setal cell development (Fig. 3a,b). We did not observe any elongated setal cells or cell files of setal cells in Mp*seta*^*ko*^ mutants. The SF/SP ratio (a ratio of the length from the foot to the proximal side of sporangium to the total sporophyte length) and the number of cells around the seta region were significantly reduced in Mp*seta*^*ko*^ compared with the wild type (Fig. 3c,d). A detailed analysis of earlier stages of sporophyte development revealed that defects in Mp*seta-1*^*ko*^ setae could be found even at stage VI, the earliest stage at which putative seta mother cells are unequivocally recognized (Fig. 3e). Despite the obvious loss of setae, spores and other sporophytic tissues were normally formed in Mp*seta-1*^*ko*^ (Fig. 3e,f). We concluded that Mp*seta*^*ko*^ mutants have defects in the differentiation from putative seta precursor cells to seta mother cells, which prevents the induction of subsequent proliferative cell divisions. At one-month postfertilization, the wild-type sporangia were pushed out of the calyptras. In contrast, Mp*seta*^*ko*^ sporangia remained buried inside the calyptras, presumably due to the defects in seta development, and hence were not exposed to the outside (Fig. 3g).

**Fig. 3.**
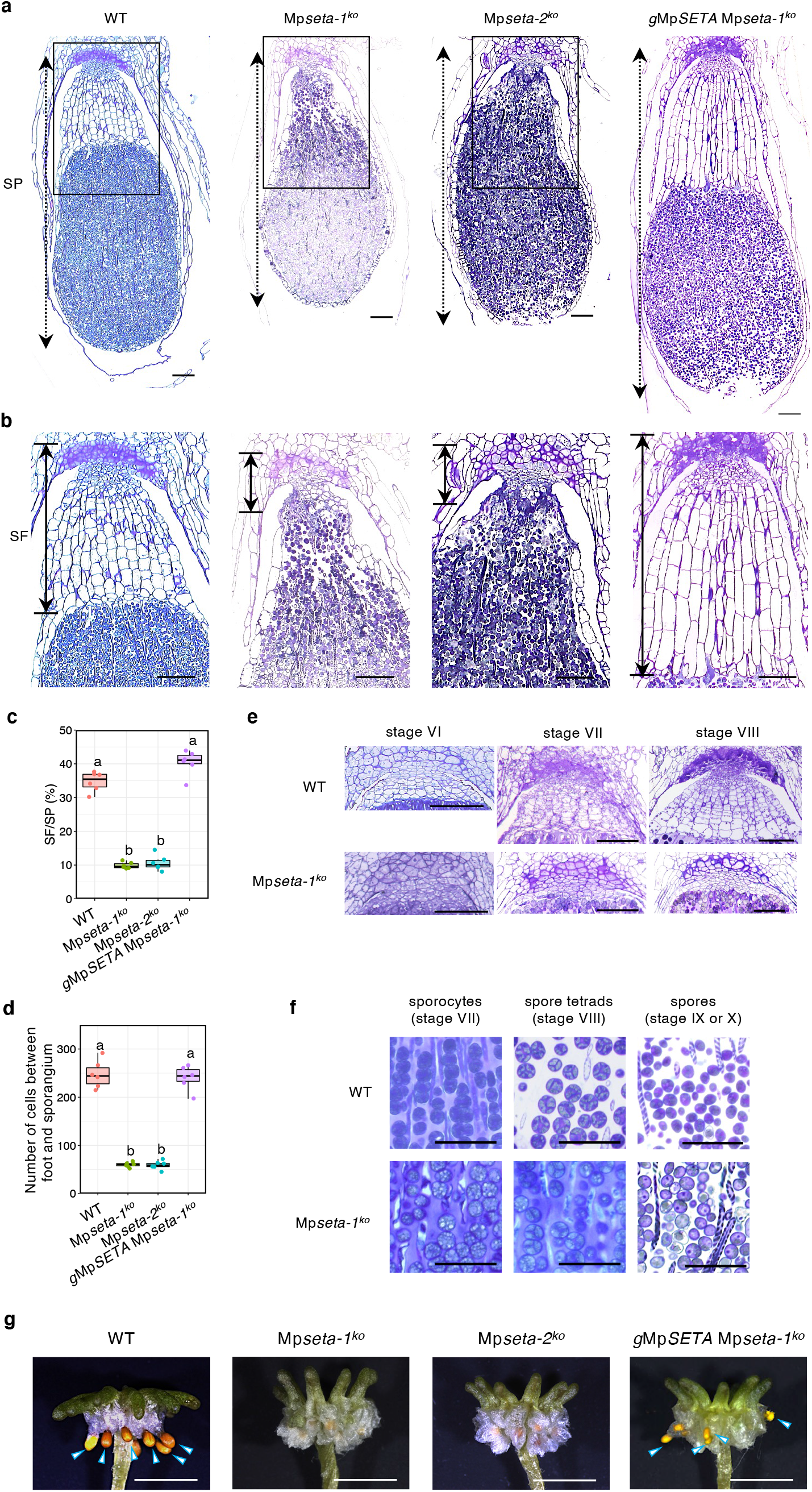
Mp*seta*^*ko*^ mutants show developmental defects in the seta. **a**,**b**, Longitudinal sections of sporophytes of the indicated genotypes. Enlarged images of the square areas are shown in (**b**). Dashed and solid arrows indicate the length of the sporophyte (SP) and the length from the boundary between the gametophyte and sporophyte to the proximal side of the sporangium (SF), respectively. **c**,**d**, Quantitative data showing the SF/SP length ratio and the number of cells in the seta region. Since the boundary between the seta and foot is unclear, cells excluding transfer cells (the cells at the boundary between gametophyte and sporophyte) were used to count the number of cells. The different letters indicate significantly different mean values at *p* < 0.01 (Tukey’s HSD test) (*n* = 6 independent lines). **e**, Comparative analyses of the tissue development between wild type (WT) and Mp*seta-1*^*ko*^. Enlarged images of longitudinal sections of stage VI, stage VII, and stage VIII sporophytes. **f**, Sporogenesis process in wild type (WT) and Mp*seta-1*^*ko*^. Sporocytes are the cells before the pre-meiotic stage, and spore tetrads are the cells immediately after meiosis. **g**, One month-postfertilization archegoniophores that produce mature sporophytes. Arrowheads indicate sporangia exposed to the outside of the calyptra due to the elongation of seta cells. Bars, 50 μm (**f**), 100 μm (**a, b, e**), and 5 mm (**e**).

Because sporangia are completely wrapped by calyptras and pseudoperianths, spore dispersal does not occur unless the sporangia are pushed out of the calyptras by seta cell elongation^29^. In genetic complementation experiments, we generated a transgenic line with the genomic region of Mp*SETA* introduced into Mp*seta-1*^*ko*^ (*g*Mp*SETA* Mp*seta-1*^*ko*^). The sporophytes obtained by crossing these complemented lines had normal setae, and their sporangia were pushed out of the calyptra as in the wild type (Fig. 3). Consequently, these results suggest that Mp*SETA* is essential for the setal formation and spore dispersal in *M. polymorpha*.

### MpICE2 physically interacts with MpSETA and regulates setal development

Because the interaction between Ia and IIIb bHLHs is evolutionarily conserved^9,10,30^, we examined whether Ia bHLH and IIIb bHLH physically interact with each other in *M. polymorpha*. We performed phylogenetic analyses to find IIIb bHLH genes in *M. polymorpha* and identified two genes encoding IIIb bHLH proteins in the genome of *M. polymorpha*, both of which are orthologous to At*ICE1* and At*SCRM2* (Extended Data Fig. 5). We named these genes Mp*ICE1* (Mp4g04910) and Mp*ICE2* (Mp4g04920). The amino acid sequences of the bHLH and ACT-like domains of IIIb bHLHs were highly conserved among land plants (Extended Data Fig. 6). Additionally, two putative IIIb bHLH-encoding genes, Lc*ICE1* and Lc*ICE2*, were found in the genome of *L. cruciata* (Extended Data Figs. 5 and 6).

Reanalyzing public RNA-seq data revealed high expression of Mp*ICE2* at 13 days postfertilization in young sporophytes (from stage III to stage IV)^19^, in which putative seta mother cells were dividing and differentiating, whereas the expression level of Mp*ICE1* was almost constant in all tissues (Fig. 4a). Thus, we assumed that MpICE2 may function predominantly in cooperation with MpSETA during seta cell formation rather than MpICE1. We expressed *Citrine* (a yellow fluorescent protein; YFP), *GUS*, and the nuclear localization-signal (*NLS*) fusion gene (*Citrine-GUS-NLS*) under the control of the Mp*ICE2* promoter to confirm the tissue/cell-level expression pattern of Mp*ICE2* in the sporophyte. In this line, GUS signals were detected overall in stages IV and V sporophytes, specifically in the seta and foot in stages VI–VIII sporophytes, and only in the foot in mature sporophytes together with Citrine and GUS signals in gametophytic tissues (Extended Data Fig. 7).

**Fig. 4.**
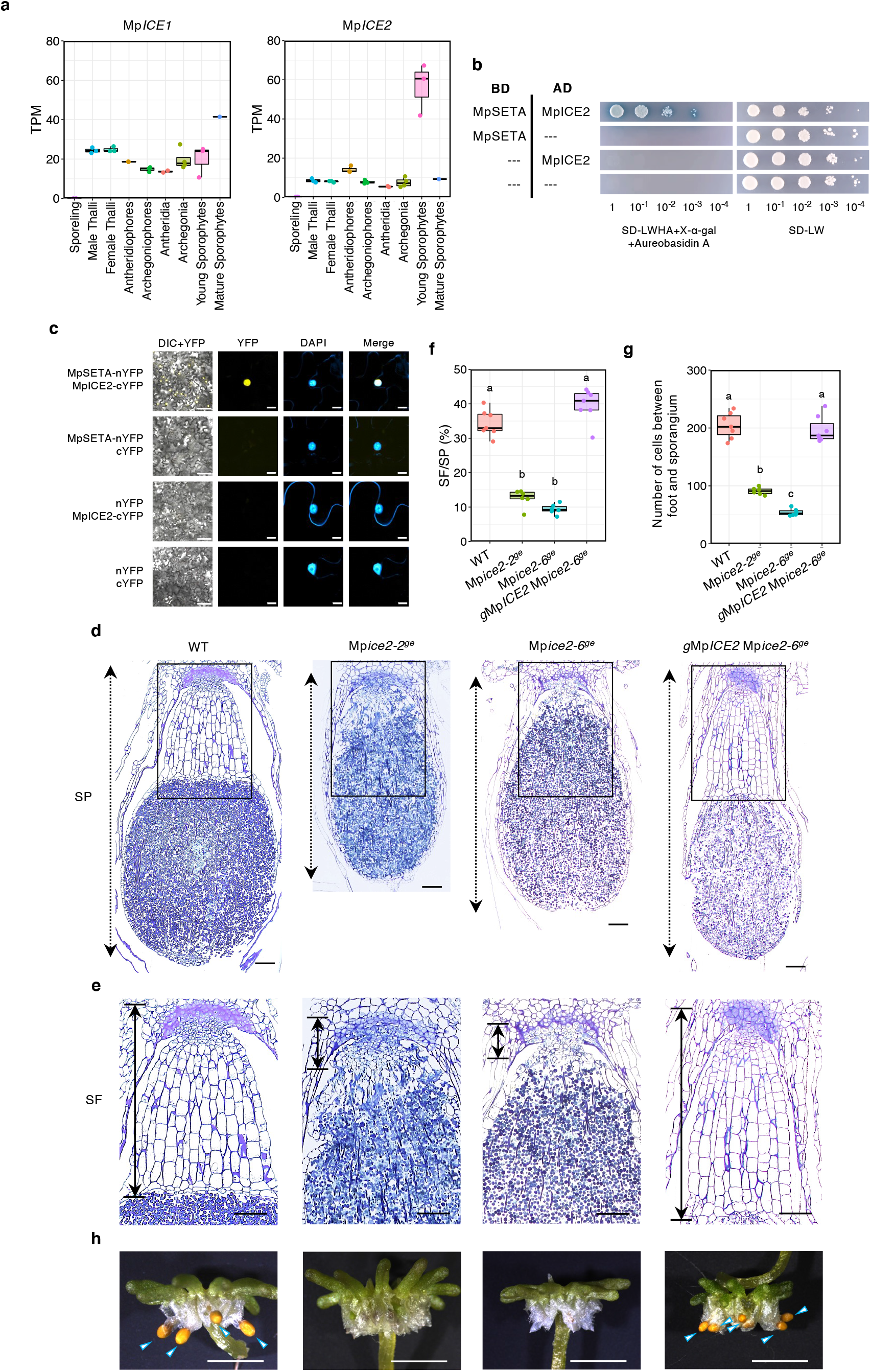
MpICE2 regulates setal development in cooperation with MpSETA. **a**. Box plot showing the expression profiles of Mp*ICE1* and Mp*ICE2* across nine *M. polymorpha* tissues. The Y-axis shows transcripts per million (TPM). **b**, Y2H assays showing the interaction between MpSETA and MpICE2. MpSETA fused with the GAL4 DNA-binding domain (DBD) was used as bait, whereas MpICE2 fused with the GAL4 activation domain (AD) was used as prey. DBD alone and AD alone were used as negative controls. **c**, BiFC assays showing the interaction between MpSETA and MpICE2 in *N. benthamiana* leaf epidermal cells. MpSETA was fused to the N-terminal fragment of EYFP (nYFP), whereas MpICE2 was fused to the C-terminal fragment of EYFP (cYFP). nYFP alone and cYFP alone were used as the negative controls. Nuclei were stained by 4′,6-diamidino-2-phenylindole (DAPI). **d**,**e**, Longitudinal sections of sporophytes of indicated genotypes. Enlarged images of the square areas are shown in (**e**). Dashed and solid arrows indicate the length of the sporophyte (SP) and the length from the boundary between the gametophyte and sporophyte to the proximal side of the sporangium (SF), respectively. **f**,**g**, Quantitative data showing the SF/SP length ratio and the number of cells in the seta region. Because the boundary between the seta and foot is unclear, cells excluding transfer cells (the cells at the boundary between gametophyte and sporophyte) were used to count the number of cells. The different letters indicate significantly different mean values at *p* < 0.01 (Tukey’s HSD test) (*n* = 6 or 7 independent lines). **h**, One month-postfertilization archegoniophores that produce mature sporophytes. Arrowheads indicate sporangia exposed to the outside of the calyptra due to the elongation of seta cells. Bars, 10 μm (**c**, YFP, DAPI, and Merge), 100 μm (**c**, left, **d**, and **e**), and 5 mm (**f**).

We performed yeast two-hybrid assays using full-length MpSETA in pairwise combinations with MpICE2 to test the interaction between MpSETA and MpICE2; in this context, a physical interaction was observed (Fig. 4b). Additionally, a BiFC assay was used to test the interaction between these bHLH TFs. YFP signals were detected in the nuclei of *Nicotiana benthamiana* leaves coexpressing MpSETA-nYFP and MpICE2-cYFP (Fig. 4c). These results suggest that MpSETA can interact with MpICE2 in *M. polymorpha*.

We generated two independent genome-edited lines using the CRISPR/Cas9 system (Mp*ice2-2*^*ge*^ and Mp*ice2-6*^*ge*^) to assess the function of Mp*ICE2*^31,32^. The first one retained two amino acid substitutions (L503H and M504L) that were predicted to be important for the DNA-binding activity of the bHLH domain^33^, whereas the second had a frame-shift mutation that caused deletion of the C-terminal half of the bHLH domain and the whole ACT-like domain (Extended Data Fig. 8). In these Mp*ice2* mutants, the setae of sporophytes were not formed (Fig. 4d,e), and the number of cells between the foot and sporangium was significantly reduced (Fig. 4f,g). Additionally, we found that the number of cells in the seta region of Mp*ice2-2*^*ge*^ exceeded that of Mp*ice2-6*^*ge*^ (Fig. 4g). This is probably because the predicted translational product of Mp*ice2-2*^*ge*^ has a two-amino acid substitution in the bHLH domain and might be partially functional in comparison with the null allele Mp*ice2-6*^*ge*^. In both Mp*ice2*^*ge*^ mutants, the sporophytes did not emerge outside of the protective organs derived from the archegonia, similar to the Mp*seta*^*ko*^ mutants (Fig. 4h). Since the single mutants of Mp*ice2*^*ge*^ showed almost the same phenotype as that of the Mp*seta*^*ko*^ mutants, Mp*ICE1* may not be functionally redundant with Mp*ICE2*, at least in setal formation. The Mp*ice2*^*ge*^ phenotype in the setal region was completely suppressed by introducing the genomic region of Mp*ICE2* into Mp*ice2-6*^*ge*^ (Fig. 4d-h). Therefore, we can conclude that the MpSETA-MpICE2 heterodimer plays an important role in the setal development of *M. polymorpha*.

Next, we tested if Mp*ICE2* can enhance the rescue of the stomatal phenotype of *mute-2* by _*pro*_At*MUTE:*Mp*SETA*. The overexpression of Mp*ICE2* in _*pro*_At*MUTE:*Mp*SETA mute-2* did not enhance stomatal formation (Extended Data Fig. 9a) (3.63 ± 3.16 and 2.64 ± 1.41 per abaxial side of the cotyledon in the lines #8-4 and #10-11, respectively [mean ± s.d.; *n* = 11]). Therefore, the MpSETA-MpICE2 heterodimer does not appear to regulate the expression of stomatal genes in *A. thaliana*. Additionally, the expression of Mp*ICE1* or Mp*ICE2* under the control of the At*ICE1* promoter in *ice1-2 scrm2-1* mutants failed to cause stomatal-lineage cell formation (Extended Data Fig. 9b). Therefore, MpICE1 and MpICE2 cannot act with AtSPCH to regulate stomatal formation in *A. thaliana*, despite the similarity of their amino acid sequences with AtICE1 (Extended Data Fig. 6).

## Discussion

In this article, we showed that two transcription factors, MpSETA (Ia bHLH) and MpICE2 (IIIb bHLH), play a pivotal role in controlling the formation of the diploid tissue seta in the sporophyte of *M. polymorpha*, which is an astomatous liverwort (Figs. 3 and 4). Similarly, in other non-liverwort land plants, a module formed by subfamilies Ia and IIIb bHLH TFs regulates GRNs in stomatal development^2^. Mp*SETA* could partially complement the defects of *A. thaliana mute* and *fama*, suggesting similar properties of Ia bHLH TFs from *M. polymorpha* and *A. thaliana*. However, Mp*SETA* was unable to complement the *spch* mutant (Fig. 1b and Extended Data Fig. 2). These results corroborate the previous hypothesis, stating that the ancestral Ia bHLH proteins had a MUTE- and FAMA-like function^34,35^. Although the nature of Mp*SETA-*expressing cells remains unknown, Mp*SETA* may function as a regulator of cell differentiation and asymmetric cell division during the setal formation, similar to its role in stomatal formation (Fig. 5a).

**Fig. 5.**
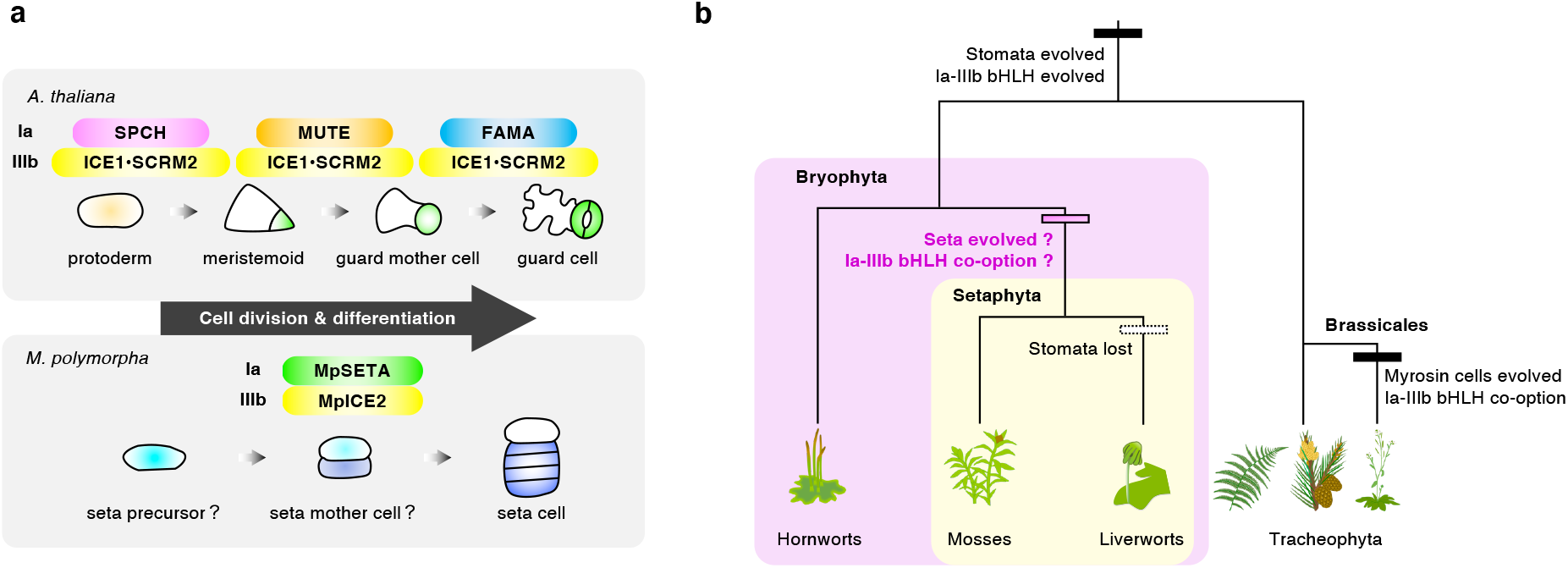
Function and co-option of the Ia-IIIb bHLH module during land plant evolution. **a**, Schematic model comparing the molecular functions of the Ia-IIIb bHLH TF modules in *A. thaliana* and *M. polymorpha* during cell fate determination. In *A. thaliana*, the heterodimers of Ia bHLHs (SPCH, MUTE, and FAMA) and IIIb bHLHs (ICE1 and SCRM2) control stomatal development. In *M. polymorpha*, the heterodimer of Ia bHLH (MpSETA) and IIIb bHLH (MpICE2) regulates cell differentiation and cell division in the seta precursor transition and might be directly or indirectly involved in the symmetric division of a putative seta mother cell. **b**, An evolutionary model for the Ia-IIIb bHLH TF module. The co-option of the Ia-IIIb bHLH module might have occurred multiple times independently during land plant evolution. First, stomata and a transcriptional module comprising Ia-IIIb bHLHs evolved in the common ancestor of land plants. In the ancestral plant, the Ia bHLHs may have had MUTE- and FAMA-like functions. Second, the Ia-IIIb bHLH TF module might have been co-opted to regulate setal development in the ancestor of the Setaphyta. Third, after mosses and liverworts diverged, the common ancestor of liverworts lost its stomata. A co-option of the Ia-IIIb bHLH module occurred in the Brassicales plants for regulating myrosin idioblast development.

Although the stomata and setae completely differ morphologically, both play a common role in promoting sporangial dehiscence and spore dispersal. In mosses and hornworts, stomata are present in the epidermis of the sporangia and function in the desiccation of the sporangia by gas exchange to promote its dehiscence and spore dispersal^25,36^. In *A. thaliana*, At*ICE1* controls stomatal development on the anther epidermis and can regulate dehydration and dehiscence of the anther^37^. However, the seta is the tissue that supports the sporangia and plays a role through cell elongation in thrusting the sporangia outside the surrounding maternal tissues to permit the long-distance dispersal of spores^24,25^. In this study, we found that the lack of setae in Mp*seta*^*ko*^ and Mp*ice2*^*ge*^ mutants prevented sporangia dehiscence and spore dispersal due to the inability of the sporangia to break through the protective organs around it (Figs. 3 and 4). Thus, the Ia-IIIb bHLH module plays a common role in the development of the tissues involved in spore or pollen dispersal.

Previous research suggested that land plant evolution occurred through the reutilization and/or modification of preexisting GRNs^20,38–43^. Here, we hypothesize that the Ia-IIIb bHLH module was primarily used in stomatal formation and secondarily co-opted to setal formation in the common ancestor of “Setaphyta” (Fig. 5b). Because only mosses and liverworts have setae, a moss–liverwort clade is called Setaphyta^1,44,45^. However, the process of setal development differs between mosses and liverworts; whereas a transient intercalary meristem, called the seta meristem, produces seta in mosses, the body plan, including seta, foot, and sporangium, is established by the formative cell division at an early stage in liverworts^25^. Therefore, whether the Ia-IIIb bHLH module is involved in the formation of seta in mosses, such as *P. patens*^46,47^, should be tested.

In *A. thaliana*, the AtFAMA-AtICE1 heterodimer regulates not only the development of stomata but also of myrosin idioblasts, which are adjacent to vascular tissues and contribute to the defense against herbivores^41,48,49^. Because myrosin cells are present only in the Brassicales, the AtFAMA-AtICE1 module for stomatal formation was co-opted for myrosin cell development during the evolution of Brassicales (Fig. 5b). The AtFAMA-AtICE1 module is thought to regulate the expression of different genes in stomatal and myrosin cell lineages^41,48,49^; however, the detailed mechanism remains unknown. Liverworts have already lost several stomatal-related genes^4,50^, such as the leucine-rich-repeat receptor-like gene *TOO MANY MOUTHS* (*TMM*) and the secreted peptide gene *EPIDERMAL PATTERNING FACTOR 1/2* (*EPF1/2*). The mechanisms underlying the development of setal cell lineage may differ from those related to stomatal cell lineages. AtMUTE directly regulates cell cycle-related genes (cyclin and cyclin-dependent kinase genes) and several stomatal-related TF genes^51^, and AtSPCH directly regulates stomatal-related genes and brassinosteroid pathway genes^52^. In *M. polymorpha*, orthologues of many of the Ia-IIIb bHLH target genes, including *CYCB, CYCD, CDKB, ERECTA*, and BZR/BES family TF genes, are conserved^4^. RNA-seq and ChIP-seq analyses could identify genes that function downstream of the MpSETA-MpICE2 module and help to clarify whether other stomatal formation-related genes are involved in the setal formation.

Are *SMF* genes and/or Mp*SETA* orthologs conserved in the Jungermanniopsida or Haplomitriopsida liverworts? A BLAST search of 1,000 plant transcriptomes (OneKP)^53^, using AtFAMA as a query, revealed the absence of Ia bHLH TFs in liverworts except for MpSETA. Because the liverwort transcriptome samples used in OneKP often do not contain sporophytes, detecting genes specifically or transiently expressed in developing sporophytes, such as Ia bHLH, is difficult. Additionally, owing to the loss of seta in the Ricciaceae species, investigating whether these species have Ia bHLH and whether the Ia bHLHs are functional to understand seta evolution is crucial. Genome analyses of various plant species in the future could be useful for understanding the relationship between Ia bHLH diversification and stomata/seta formation.

## Methods

### Phylogenetic analysis

The bHLHs were classified according to Pires and Dolan^5^. We retrieved the amino acid sequence information from each of the following; MarpolBase (https://marchantia.info), Phytozome v.13 (https://phytozome-next.jgi.doe.gov/), OneKP (https://db.cngb.org/onekp/), TAIR (http://www.arabidopsis.org/), and NCBI (https://www.ncbi.nlm.nih.gov/genome/?term=PRJNA701193). Then, we aligned the bHLH domain and the C-terminal ACT-like domain using MAFFT^54^ v.6.864 (https://www.genome.jp/tools-bin/mafft) with the default parameters. Noting that the alignment gaps were removed manually. Next, we visualized the amino acid sequences with Jalview^55^ v.2.11.2.1. To predict the importance of the amino acids in nucleotide binding and dimer formation in the bHLH domains, we used BLAST searches of NCBI (https://blast.ncbi.nlm.nih.gov/Blast.cgi). Alternately, we performed the phylogenetic tree constructions using the maximum-likelihood algorithm on MEGA 7^56^ with the JTT+G+I and LG+G+I substitution models for Ia and IIIb bHLH, respectively. To assess the statistical support for the topology, we performed bootstrap analyses with 1,000 replicates for each analysis. Subfamilies III(a+c) and III(d+e) bHLHs were chosen as the outgroup of the phylogeny for Ia and IIIb bHLH, respectively.

### Plant materials and growth conditions

We asexually maintained the male and female accessions of *M. polymorpha* L., Takaragaike-1 (Tak-1), and Tak-2, respectively. A female progeny backcrossed to Tak-1 for three backcross generations (BC3-38) was also used as the wild type. We cultured gemmae and thalli on half-strength Gamborg’s B5 media containing 1% (w/v) agar and 1% (w/v) sucrose under 50–60 μmol photons m^−2^ s^−1^ continuous white light at 22?. To induce gametangiophore development, we transferred gemmae to 16-h-light/8-h-dark conditions with 50–60 μmol photons m^−2^ s^−1^ white light and 50–60 μmol photons m^−2^ s^−1^ far-red light emitted from diodes (IR LED STICK 18W, Namoto) at 18? and incubated for one month.

We used the *Arabidopsis thaliana* Columbia-0 (Col-0) accession as wild type except for *mute-2*, where Wassilewskija-4 (Ws-4) was used. The seeds were surface sterilized with 70% ethanol, then sown onto half-strength MS media containing 0.5% (w/v) gellan gum and 1% (w/v) sucrose. We incubated the seeds at 22°C under 50–60 μmol photons m^−2^ s^−1^ continuous white light.

### Complementation tests of *A. thaliana* stomatal defective mutants

We obtained T-DNA insertion mutants *spch-3* (SAIL_36_B06) and *fama-1* (SALK_100073) from the Arabidopsis Biological Resource Center (ABRC), while *mute-2* (FLAG_225D03) was from the French National Institute for Agricultural Research (INRA). *ice1-2 scrm2-1*^9^ was provided by K.U. Torii. To construct _*pro*_At*SPCH:*Mp*SETA*, firstly, we amplified the At*SPCH* promoter (2,572 bp upstream of the translational initiation site) and Mp*SETA* CDS from Col-0 gDNA and cDNA derived from Tak-1 antheridiophores, respectively. Then, we fused _*pro*_At*SPCH* and Mp*SETA* CDS fragments by PCR using PrimeSTAR GXL polymerase (Takara Bio). The resultant PCR fragment was subcloned into pENTR1A entry clones (Invitrogen) at the SalI and EcoRV restriction sites using an In-Fusion HD Cloning Kit (Takara Bio), which we transferred into the destination vector pGWB501^57^ using Gateway LR Clonase II Enzyme mix (Thermo Fisher Scientific). We generated the _*pro*_At*MUTE:*Mp*SETA* and _*pro*_At*FAMA:*Mp*SETA* constructs by LR recombination of the R4pGWB501^58^ or R4pGWB601^58^ with a pENTR1A containing the Mp*SETA* CDS at the EcoRI sites and either pENTR5’/TOPO_proAtMUTE harboring _*pro*_At*MUTE*, 1,930 bp upstream of the translational initiation site, or pENTR5’/TOPO_proAtFAMA harboring _*pro*_At*FAMA*, 3,105 bp upstream of the translational initiation site (R4pGWB501_proAtMUTE:MpSETA and R4pGWB601_proAtFAMA:MpSETA). To construct _*pro*_At*ICE1:*Mp*ICE1* or _*pro*_At*ICE1:*Mp*ICE2*, we amplified, respectively, the At*ICE1* promoter (2,578 bp upstream of the translational initiation site) from Col-0 gDNA, and Mp*ICE1* or Mp*ICE2* CDS from cDNA derived from Tak-1 thalli. We fused _*pro*_At*ICE1* and Mp*ICE1* or Mp*ICE2* CDS fragments by PCR using PrimeSTAR GXL polymerase. We subcloned the resultant PCR fragment into pENTR1A entry clones at the SalI and EcoRI restriction sites using an In-Fusion HD Cloning Kit and transferred them into the destination vector pGWB501^57^ using Gateway LR Clonase II Enzyme mix. We introduced the resultant plasmids into *spch-3/+, mute-2/+, fama-1/+*, or *ice1-2/+ scrm2-1* heterozygous plants by the previously described method^59^ using *Agrobacterium tumefaciens* strain GV3101. We confirmed that all the transformants had a single insertion event by segregation analyses. We used T_3_ or T_4_ homozygous plants. For Mp*ICE2* overexpression analyses, we transferred the Mp*ICE2* CDS into the destination vector pFAST-R02^60^ using Gateway Clonase II Enzyme mix, and the resultant plasmid was introduced into Ws-4, *mute-2/+*, and _*pro*_At*MUTE:*Mp*SETA mute-2/+* (#8-4 and #10-11). T_1_ seeds expressing TagRFP were selected, and T_1_ plants were used for the analyses. We stained the cotyledons with FM4-64 and observed them using an LSM780 laser scanning microscope (Carl Zeiss). Images were processed with Fiji (NIH). The sequences of the primers used in this study are shown in Supplementary Tables 1 and 2.

### Gene expression analysis

Publicly available transcriptome data were downloaded from the Sequence Read Archive (SRA) repository. Accession numbers included; sporelings (SRR4450254, SRR4450255, SRR4450256)^4^, male thalli (DRR118949, DRR118950, DRR118951)^20^, female thalli (DRR118943, DRR118944, DRR118945)^20^, antheridiophores (DRR050346, DRR050347, DRR050348)^21^, archegoniophores (DRR050351, DRR050352, DRR050353)^21^, antheridia (DRR050349, DRR050350)^21^, archegonia (DRR209029, DRR209030, DRR209031, DRR209031)^22^, 13 DPF sporophytes (SRR1553297, SRR1553298, SRR1553299)^19^, and mature sporophytes (SRR896223)^4^. We pre-processed the RNA-seq data to filter out low-quality sequences using fastp^61^ v.0.20.0. The sequence reads were mapped to the *M. polymorpha* genome v.6.1 (https://marchantia.info) by STAR^62^ v.2.7.8a with default parameters. We performed the post-processing of SAM/BAM files using SAMtools^63^ v.1.11. To calculate the transcript per million (TPM) via the RSEM^64^ v.1.3.0 with default parameters, we used the read for each gene. We created the plots using Rstudio v.1.4.1106 (https://www.rstudio.com/).

### Histology

We stained the sporophytes (stage I and II) with 4′,6-diamidino-2-phenylindol (DAPI) as described previously^22^ and observed them using an LSM780 laser scanning microscope.

We fixed the plant samples with 2% (w/v) paraformaldehyde and 2% (v/v) glutaraldehyde in 0.05 M cacodylate buffer (pH 7.4) for 2 h at room temperature, post-fixed with 2% (v/v) osmium tetroxide in 0.1 M cacodylate buffer for another 2 h at room temperature, dehydrated in an ethanol series, substituted with acetone, and then embedded in Spurr’s resin (Polysciences). We cut the Spurr’s blocks into semi-thin sections (0.75–2 μm) with glass knives on an ultramicrotome Leica Ultracut UCT (Leica Microsystems) stained with a solution containing 1% (w/v) sodium tetra-borate and 1% (w/v) toluidine blue O. The sections were mounted on MAS-coated glass slides (Matsunami Glass). In the end, we obtained the images using a VB-7010 (KEYENCE)/AxioCam HRc (Zeiss) and processed them with either Fiji or Adobe Photoshop Elements 9 (Adobe Systems).

### Histochemical GUS staining

To construct _*pro*_Mp*SETA:GUS*, we amplified the genomic fragment of the 4,194 bp upstream region of the translational initiation site from Tak-1 gDNA using PrimeSTAR Max DNA polymerase (Takara Bio), subcloned it into pENTR1A at the EcoRI restriction sites using the In-Fusion HD Cloning Kit and then transferred it into the destination vector pMpGWB104^65^ using Gateway LR Clonase II Enzyme mix (Thermo Fisher Scientific). To construct _*pro*_Mp*ICE2:Citrine-GUS-NLS*, firstly we amplified the genomic fragment containing the 3,060 bp upstream region of the translational initiation site from Tak-1 gDNA, *Citrine* ORF from pMpGWB107^65^, and *GUS-NLS* ORF from pPZP211_35S-NG-GUS-NLS-nosT^66^. These fragments were fused by PCR, subcloned into pENTR1A at the SalI and EcoRV sites using In-Fusion HD Cloning Kit, and transferred into pMpGWB101^65^. We introduced the resultant plasmids into Tak-1 accession by the previously described method^67^ using *A. tumefaciens* strain GV2260.

The tissues of _*pro*_Mp*SETA:GUS* or _*pro*_Mp*ICE2:Citrine-GUS-NLS* plants were vacuum-infiltrated and incubated at 37? overnight in GUS staining solution containing 10 mM EDTA (pH 8.0), 100 mM NaH_2_PO_4_ (pH 7.0), 0.1% (v/v) Triton X-100,0.5 g L^−1^ 5-bromo-4-chloro-3-indolyl-*β*-D-glucuronic acid (X-Gluc), 5 mM potassium-ferrocyanide, and 5 mM potassium-ferricyanide. We washed the samples in 70% (v/v) ethanol and cleared them with chloral-hydrate / glycerol solution.

### Generation of Mp*seta*^*ko*^ mutants

We amplified the Tak-1 genomic sequences 3,125 bp upstream and 3,101 bp downstream of the MpSETA bHLH domain coding region by PCR with a PrimeSTAR Max DNA polymerase and inserted them into the PacI and AscI sites of pJHY-TMp1^28^, respectively. We introduced the vector into germinating F_1_ spores from Tak-1 and Tak-2 crosses via *A. tumefaciens* strain GV2260 as previously described^68^. We selected the transformed T_1_ plants carrying the targeted insertion by PCR using GoTaq DNA polymerase (Promega). As T_1_ plants of Mp*seta-1*^*ko*^ and Mp*seta-2*^*ko*^ were both females, we obtained the male mutants from F_1_ sporelings by crossing female mutants with Tak-1.

### Reverse transcription PCR

For gene expression analysis, we collected 21 DPF sporophytes from wild-type, Mp*seta-1*^*ko*^, and Mp*seta-2*^*ko*^. We extracted the total RNA by RNeasy Plant Mini Kit (Qiagen) according to the manufacturer’s protocol. We evaluated total RNA qualitatively and quantitatively using a NanoDrop 2000 spectrophotometer (Thermo Fisher Scientific). First-strand cDNA was synthesized using ReverTra Ace qPCR RT Master Mix with gDNA Remover (Toyobo), and semiquantitative RT-PCR was undertaken using Mp*EF1α* as a loading control^69^.

### Yeast two-hybrid assay

We amplified the coding sequences of Mp*SETA* and Mp*ICE2* from cDNA derived from mRNA of Tak-1 thalli by PCR using PrimeSTAR Max DNA polymerase or PrimeSTAR GXL polymerase (Takara Bio). Also, we amplified the coding sequences of At*ICE1* and At*SCRM2* from cDNA derived from Col-0 leaves using PrimeSTAR GXL polymerase. We subcloned the resultant PCR fragments into pENTR1A at the EcoRI sites or SalI and EcoRV sites using an In-Fusion HD Cloning Kit. To generate a bait destination vector, pDEST-GBKT7-Amp^r^, *Amp*^*r*^ was amplified from pDEST-GADT7^70^ by PCR using PrimeSTAR Max DNA polymerase, and the fragment was cloned into the SfoI site of pDEST-GBKT7^70^. The inserted fragments, MpSETA, MpICE2, AtICE1, and AtSCRM2, were transferred into pDEST-GADT7 and/or pDEST-GBKT7-Amp^r^ using Gateway LR Clonase II Enzyme mix. Bait and prey constructs were co-transformed into the yeast strain Y2HGold (Clontech) using the Frozen-EZ Yeast Transformation II Kit (Zymo Research) and the transformants were grown on the solid SD media lacking Leu and Trp (SD-LW). To examine the interaction between the bait and prey proteins, transformants were grown on the solid SD media lacking Leu, Trp, His, and adenine with 40 mg L^−1^ X-*α*-gal and 200 μg L^−1^ Aureobasidin A (SD-LWHA/X/AbA) at 30?. pDEST-GBKT7-Amp^r^ and pDEST-GADT7 were used as negative controls (empty).

### BiFC

The coding sequences of Mp*SETA*, Mp*ICE2*, At*ICE1*, and At*SCRM2* subcloned into pENTR1A, as described above, were transferred into the pB4GWnY and/or pB4GWcY/pB4cYGW binary vector^71^ using LR reaction to be fused with N-terminal or C-terminal half of YFP (Mp*SETA-nYFP*, Mp*ICE2-cYFP, cYFP-*At*ICE1*, and *cYFP-* At*SCRM2* driven by the *CaMV35S* promoter). Transformed *A. tumefaciens* strain GV3101 cells harboring expression vectors were cultured and resuspended in distilled water to a final optical density of OD_600_ = 1.0. Mixed *Agrobacterium* cultures were infiltrated into 4-week-old *Nicotiana benthamiana* leaves. The nuclei were stained with DAPI at the 1 mg L^−1^ concentration for 30 min. We observed the samples 1.5 days post-inoculation (DPI) by an LSM780 laser scanning microscope. pB4GWnY and pB4GWcY/pB4cYGW were used as negative controls (empty).

### Generation of Mp*ice2*^*ge*^ mutants

To generate Mp*ice2*^*ge*^ mutants, we edited the Mp*ICE2* locus encoding the bHLH domain using a CRISPR/Cas9 based genome-editing system, as previously described^31^. Two sgRNAs were designed to generate Mp*ice2*^*ge*^ mutants. The oligonucleotides encoding sgRNA were cloned into pMpGE_En03^31^ between BsaI sites, then introduced into pMpGE010^31^. Male and female mutants that did not harbor T-DNA containing *Cas9* were obtained from F_1_ sporelings crossed with wild type and Mp*ice2-2*^*ge*^ or Mp*ice2-6*^*ge*^.

### Complementation tests of Mp*seta*^*ko*^ and Mp*ice2*^*ge*^

To construct *g*Mp*SETA* for Mp*seta-1*^*ko*^ complementation, the genomic region containing the 4,194 bp upstream region and coding sequences was amplified from Tak-1 genomic DNA. The fragment was cloned into pENTR1A between the EcoRI sites and then introduced into pMpGWB301^65^. The resultant plasmids were introduced into female Mp*seta-1*^*ko*^. Male *g*Mp*SETA* Mp*seta-1*^*ko*^ was obtained from F_1_ sporelings produced from crosses between female *g*Mp*SETA* Mp*seta-1*^*ko*^ and male Mp*seta-1*^*ko*^. To construct *g*Mp*ICE2* for Mp*ice2-6*^*ge*^, the genomic region containing 3,060 bp upstream regions and coding sequences was amplified from Tak-1 genomic DNA. The fragment was cloned into pENTR1A between the SalI and EcoRV sites and then introduced into pMpGWB101^65^. The resultant plasmids were introduced into female Mp*ice2-6*^*ge*^. A male *g*Mp*ICE2* Mp*ice2-6*^*ge*^ was obtained from an F_1_ sporeling derived from a cross between a female *g*Mp*ICE2* Mp*ice2-6*^*ge*^ and a male Mp*ice2-6*^*ge*^.

## Acknowledgements

We thank Tsuyoshi Nakagawa (Shimane University, Japan), Shoji Mano (National Institute for Basic Biology, Japan), Shigeo S. Sugano (National Institute of Advanced Industrial Science and Technology, Japan), and Keiko U. Torii (The University of Texas at Austin, USA) for sharing the materials. We also thank Keiji Nakajima (Nara Institute of Science and Technology, Japan) for sharing the figures of plants in Fig. 5b. We are grateful to James Raymond for his critical readings of this manuscript. This work was supported by MEXT/JSPS KAKENHI grants to M.S. (JP19K06722 and JP20H05416), to K.T. (JP26711017 and JP18K06283), to Y.O. (JP18K19964), to T.M. (JP20H05905 and JP20H05906), to I.H.-N. (JP15H05776), to R.N. (JP20H04884) and T.S. (JP18K06284); Grants-in-Aid JSPS Fellows to K.C.M. (JP21J14990) and M.S. (JP12J05453) and; the Takeda Science Foundation, the Kato Memorial Bioscience Foundation, and the Ohsumi Frontier Science Foundation to M.S. J.L.-M. and Y.-T.L. were supported by Ph.D. studentships from the Darwin Trust of Edinburgh.

## Author contributions

K.C.M. and T.S. conceived and designed the research in general; K.C.M. performed most of the experiments and analyzed the data; M.S. and Y.M. performed the experiments on Mp*SETA*; J.L.-M., Y.-T. L., G.I., and J.G. performed the experiments on Mp*ICE2*; K.T., Y.O., T.M., I.H.-N., R.N., J.G., and T.K. supervised the experiments; K.C.M. and T.S. wrote the manuscript; All authors read, edited, and approved the manuscript.

## Extended Data

**Extended Data Fig. 1.**
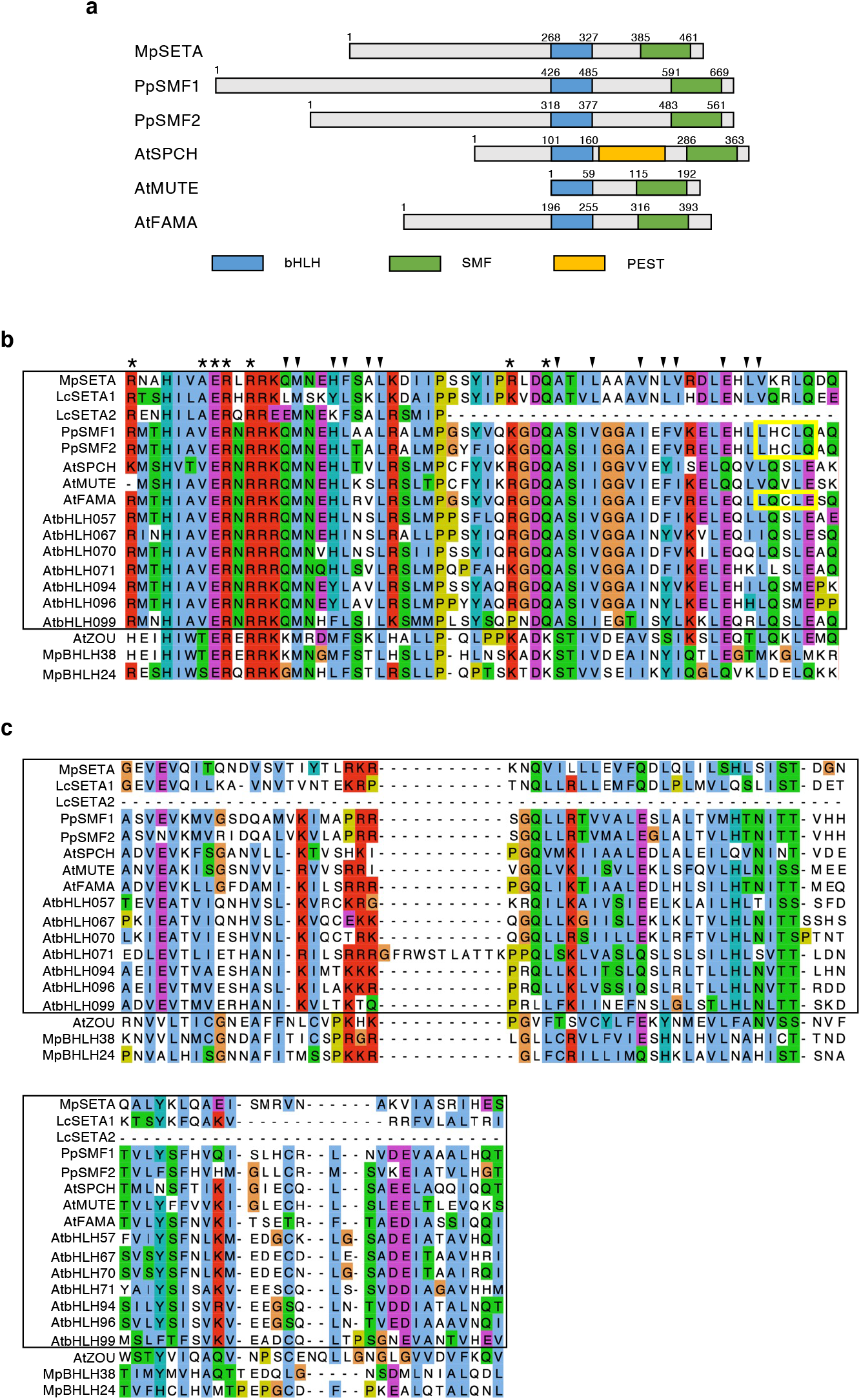
Comparison of the domain architecture of Ia bHLHs in land plants. **a**, A diagram of the domain architecture of MpSETA (*M. polymorpha*), PpSMF1, PpSMF2 (*P. patens*), AtSPCH, AtMUTE, and AtFAMA (*A. thaliana*). While no PEST domain was identified, MpSETA has a bHLH domain and SMF domain conserved at the C-terminus like other Ia bHLH proteins. SMF domain is structurally considered to be the ACT-like domain, which is a putative domain for protein-protein dimerization. **b**, Sequence alignment of the bHLH domain of Ia bHLH proteins. Ia bHLHs are surrounded by a black box, and others are Ib(1) bHLHs. Asterisks indicate amino acids that are assumed to be important for binding to the E-box (CANNTG), and the triangles indicate amino acids that are assumed to be important for the dimerization of the bHLH domain. The yellow box indicates the LxCxE motif, which is a binding motif with Retinoblastoma-related (RBR). **c**, Sequence alignment of the C-terminal SMF domain of Ia bHLH proteins.

**Extended Data Fig. 2.**
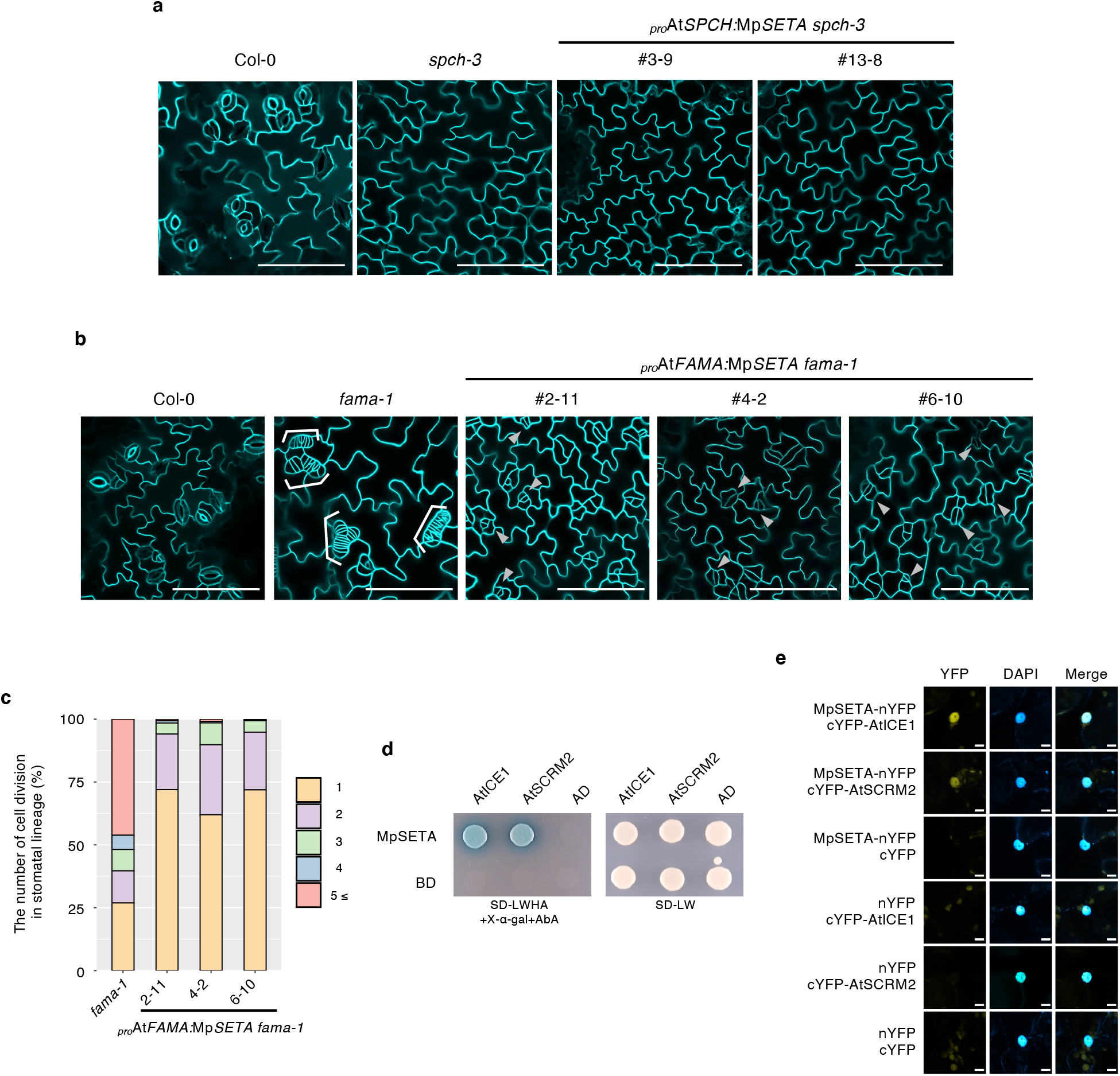
Function of Mp*SETA* in *A. thaliana* Ia bHLH mutants. **a**, Confocal images of *A. thaliana* abaxial cotyledons of wild type (Col-0), *spch-3*, and _*pro*_At*SPCH:*Mp*SETA spch-3* at 9 days after stratification (DAS). **b**, Confocal images of *A. thaliana* abaxial cotyledons of wild type (Col-0), *fama-1*, and _*pro*_At*FAMA:*Mp*SETA fama-1* at 9 DAS. Brackets and arrows indicate *fama* tumors and stomatal-lineage cells, respectively. **c**, Quantitative data of the distribution of the number of cell divisions that occurred in the stomatal lineage in each genotype. (*n* > 320 cells per genotype, 9 DAS cotyledons). **d**, Y2H assays in which the MpSETA fused with the GAL4 DNA-binding domain (DBD) was used as bait, and the AtICE1 and AtSCRM2 fused with the GAL4 activation domain (AD) were used as prey. DBD alone and AD alone were used as negative controls. **e**, BiFC assays showing the interaction between MpSETA and AtICE1 or AtSCRM2 in *N. benthamiana* leaf epidermal cells. MpSETA was fused to the N-terminal fragment of EYFP (nYFP), whereas AtICE1 or AtSCRM2 was fused to the C-terminal fragment of EYFP (cYFP). nYFP alone and cYFP alone were used as the negative controls. Nuclei were stained with DAPI. Bars, 10 μm (**e**), and 100 μm (**a**,**b**).

**Extended Data Fig. 3.**
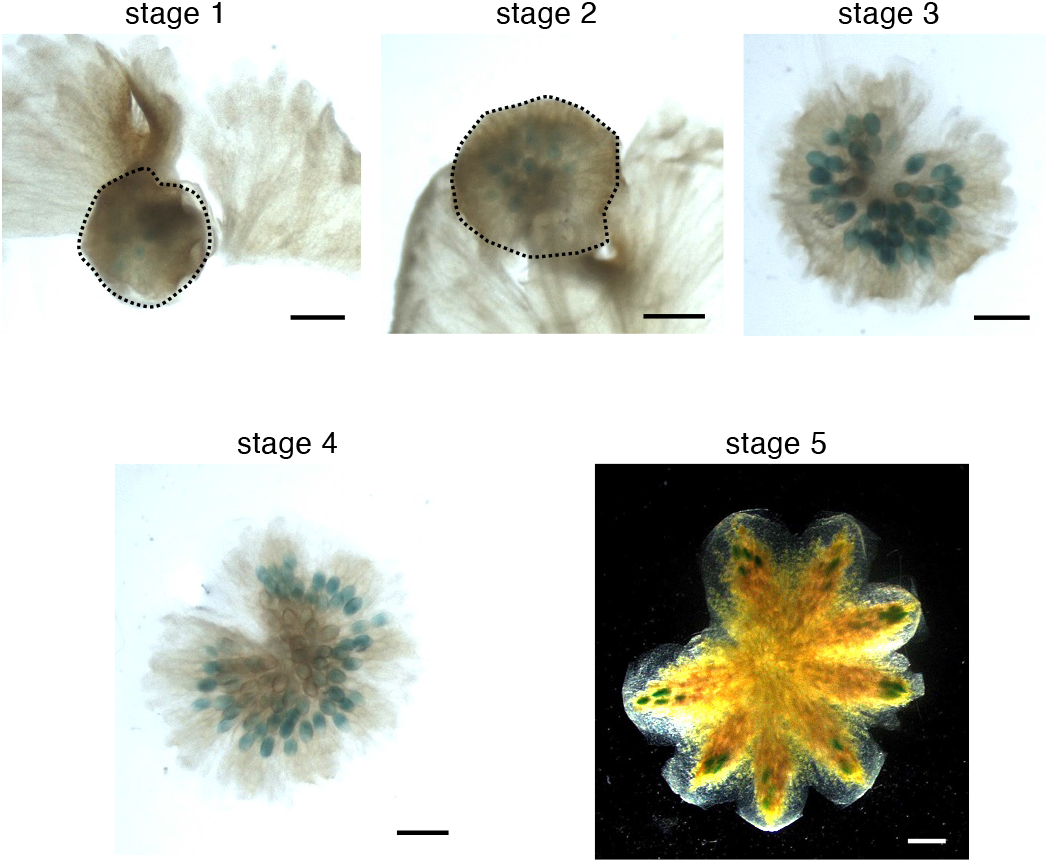
Expression analysis of Mp*SETA* in the gametophytic tissues. Histochemical detection of *β*-glucuronidase (GUS) activity driven by the Mp*SETA* promoter in the developing antheridia. Bars, 1 mm.

**Extended Data Fig. 4.**
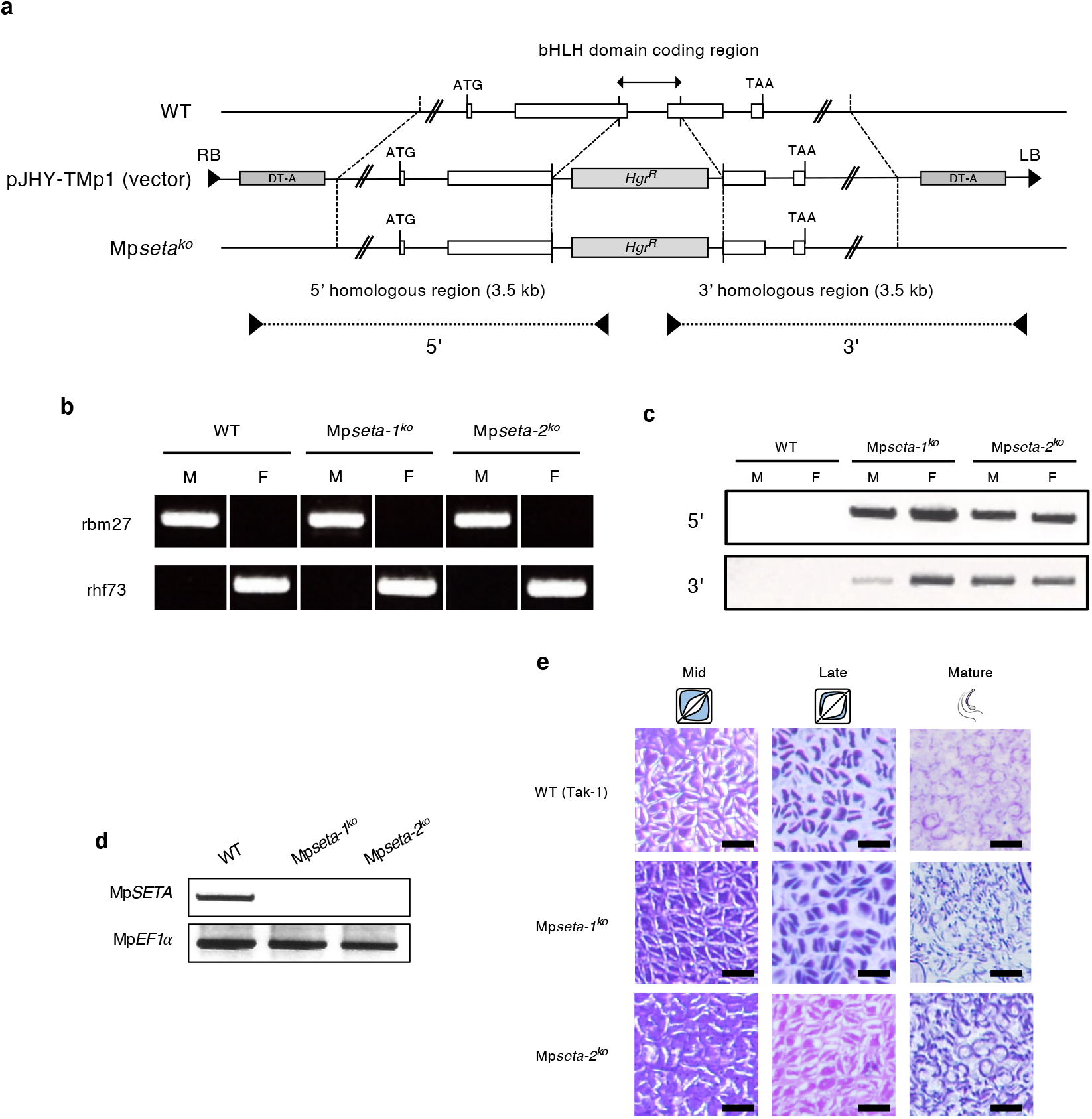
Generation and phenotypes of Mp*SETA* knock-out lines. **a**, Structure of the Mp*SETA* locus disrupted by homologous recombination. Knock-out lines have a deletion in the bHLH domain coding region. White boxes indicate the exons of the MpSETA coding sequence. DT-A, diphtheria toxin A fragment gene; *Hgr*^*R*^, hygromycin-resistance gene. **b**, Genotyping of the Mp*seta*^*ko*^ lines used in this study to distinguishsex. rbm27, a male-specific marker; rhf73, a female-specific marker. **c**, Genotyping of the Mp*seta*^*ko*^ lines. The position of the primers used for PCR is shown in (**a**). M, Male; F, Female. **d**, RT-PCR to confirm the loss of the full-length Mp*SETA* transcript in Mp*seta*^*ko*^ lines in 21 DPF sporophytes. Mp*EF1α* was used as an internal control. **e**, Spermatogenesis process in the wild type (WT) and Mp*seta*^*ko*^ lines. All the images are at the same scale. Bars, 10 μm (**e**).

**Extended Data Fig. 5.**
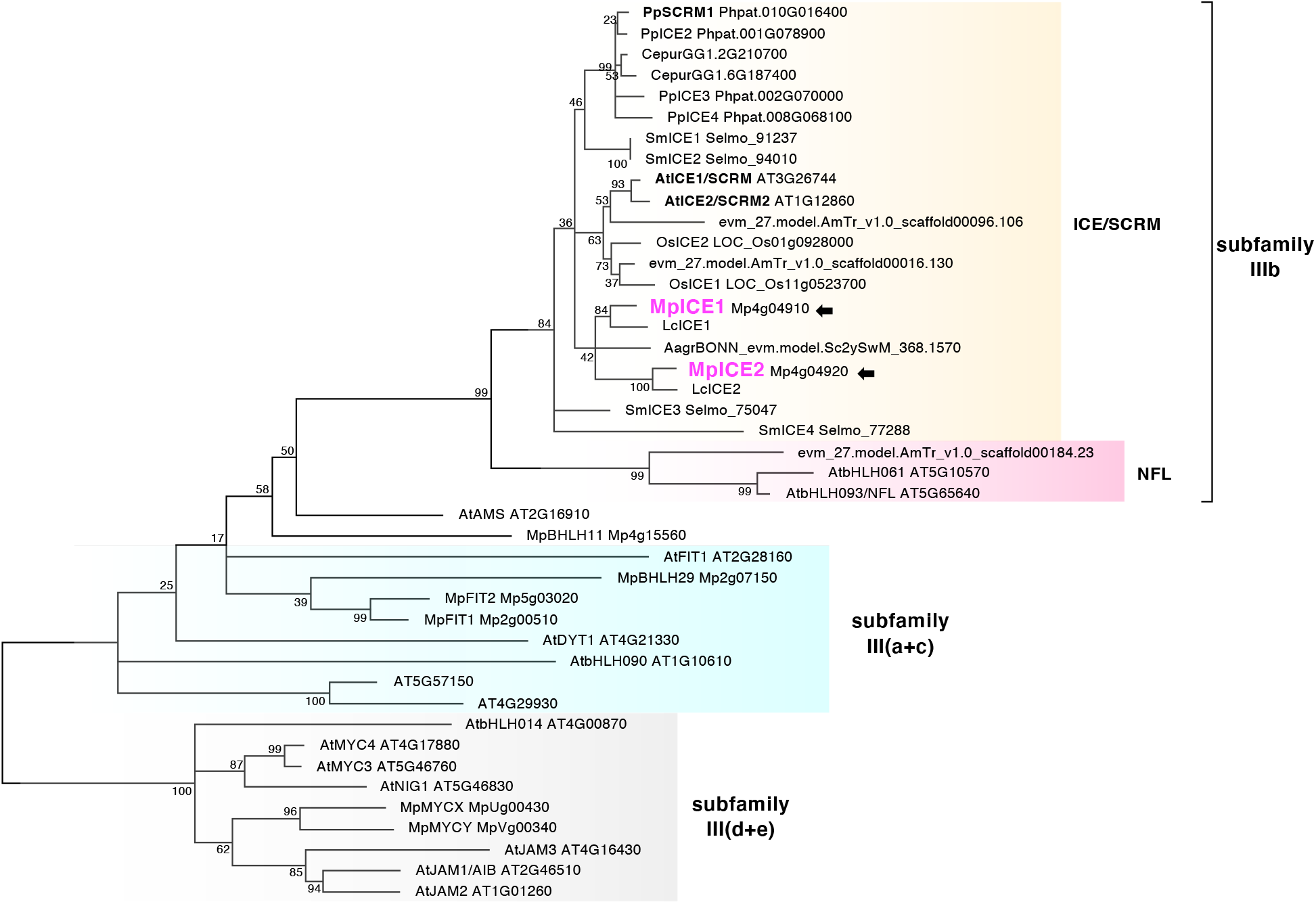
Phylogenetic tree of IIIb bHLH TFs. A maximum-likelihood bHLH phylogenetic tree of subfamilies IIIb, III (a+c) (light blue), and III(d+e) (outgroup) is shown. Numbers at branches indicate bootstrap values calculated from 1,000 replicates. IIIb bHLHs are divided into 2 groups: ICE/SCRM clade (orange) and NFL clade (magenta). Species are abbreviated as follows: Mp, *M. polymorpha* (liverwort); Lc, *L. cruciata* (liverwort); Pp, *P. patens* (moss); Cepur, *Ceratodon purpureus* (moss); Aagr, *Anthoceros agrestis* (hornwort); Sm, *Selaginella moellendorffii* (lycophyte); AmTr, *Amborella trichopoda* (basal angiosperm); Os, *Oryza sativa* (monocot); At, *A. thaliana* (dicot). Arrows indicate MpICE1 (Mp4g04910) and MpICE2 (Mp4g04920). For the phylogenetic construction of subfamilies III(a+c) and III(d+e), we used amino acid sequences from only *A. thaliana* and *M. polymorpha*.

**Extended Data Fig. 6.**
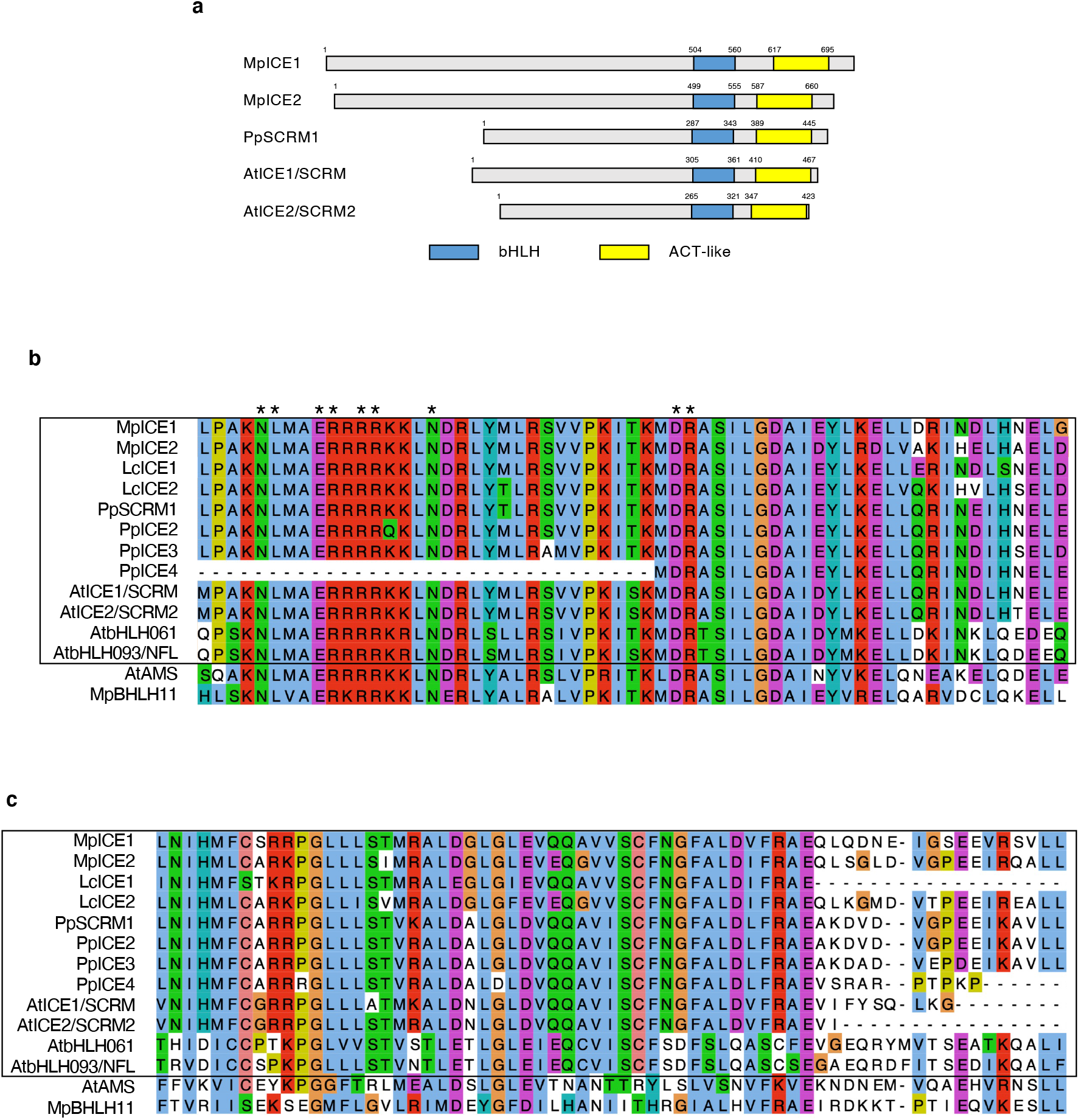
Comparison of the domain architecture of IIIb bHLHs in land plants. **a**, A diagram of the domain architecture of MpICE1, MpICE2 (*M. polymorpha*), PpSCRM1 (*P. patens*), AtICE1, and AtSCRM2 (*A. thaliana*). MpICE1 and MpICE2 have a bHLH domain and ACT-like domain conserved at the C-terminus like other IIIb bHLH proteins. **b**, Sequence alignment of the bHLH domain of the IIIb bHLH proteins. IIIb bHLHs are surrounded by a black box, and others are an outgroup. Asterisks indicate amino acids that are assumed to be important for binding to the E-box (CANNTG). **c**, Sequence alignment of the C-terminal ACT-like domain of the IIIb bHLH proteins.

**Extended Data Fig. 7.**
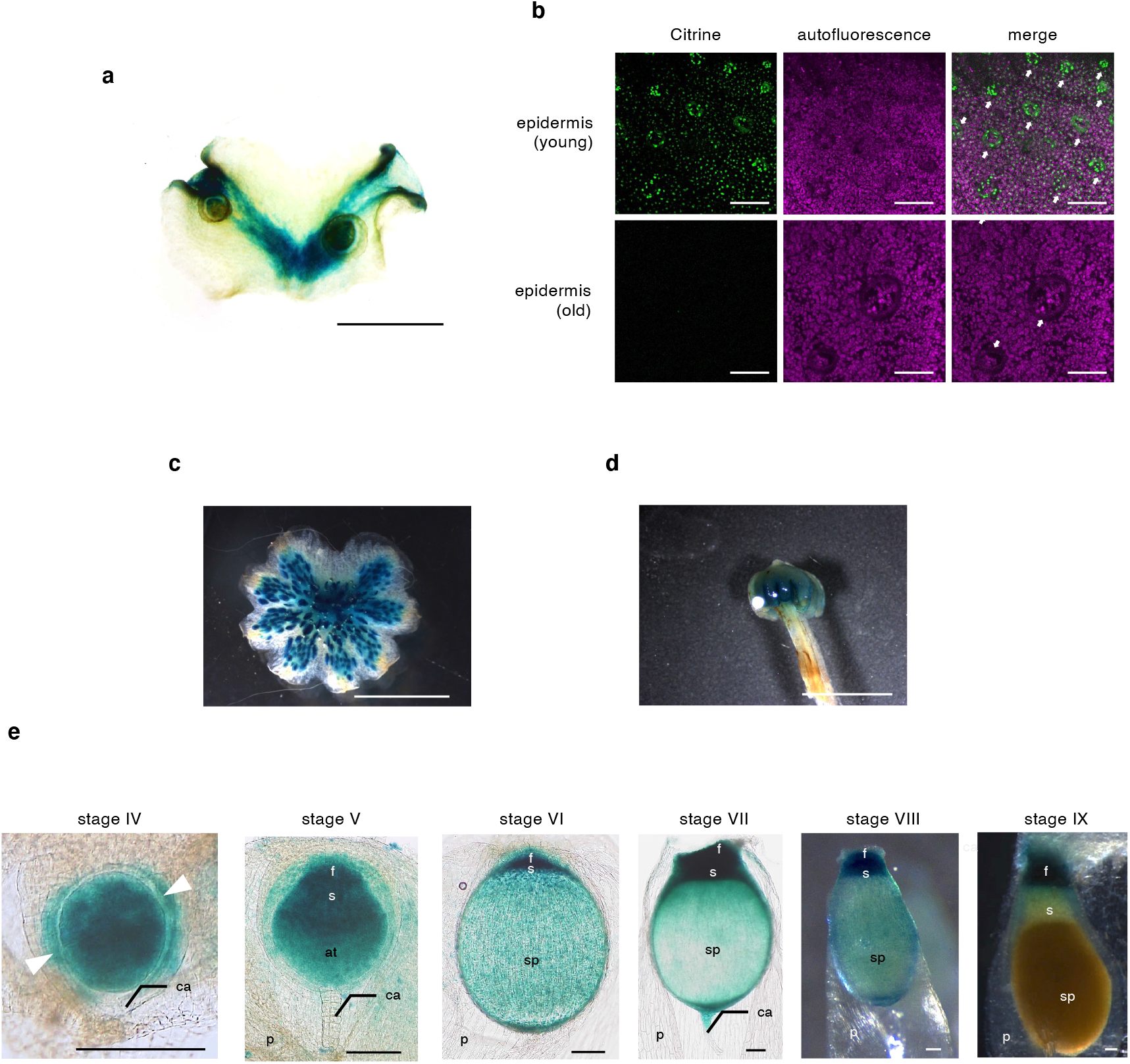
The expression analysis of Mp*ICE2*. **a**, Histochemical detection of *β*-glucuronidase (GUS) activity driven by the Mp*ICE2* promoter in the vegetative thallus. **b**, Confocal images of the dorsal epidermis of _*pro*_Mp*ICE2:Citrine-GUS-NLS* line. The upper and lower panels indicate the epidermis around the apical notch and the epidermis around the midrib, respectively. Arrows indicate the air pores. **c**,**d**, Histochemical detection of GUS activity driven by the Mp*ICE2* promoter in the gametophytic reproductive organs. An antheridiophore (**c**) and an archegoniophore (**d**) are shown. **e**, Expression pattern of Mp*ICE2* in developing sporophytes. f, foot; s, seta; at, archesporial tissue; sp, sporangium; ca, calyptra; p, pseudoperianth (*n*). Arrowheads indicate the cell wall of the first cell division. Bars, 5 mm (**c** and **d**), 100 μm (**b** and **e**).

**Extended Data Fig. 8.**
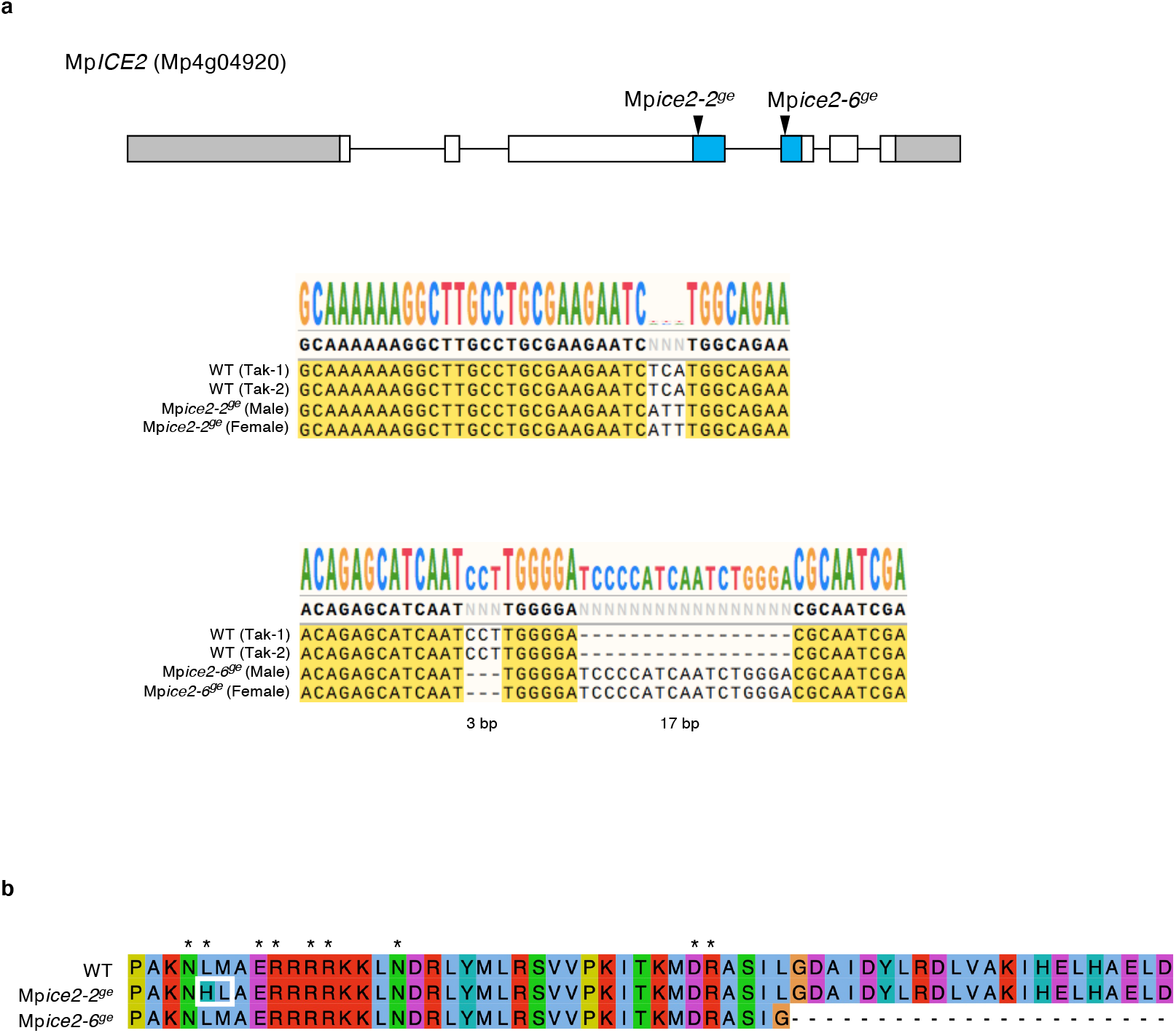
Generation of Mp*ice2* mutants by CRISPR/Cas9. **a**, Schematic representation of the Mp*ICE2* gene and the resulting mutations in the obtained CRISPR/Cas9-generated alleles. Gray, white, and blue boxes indicate the coding sequences (CDS), the untranslated regions (UTR), and the bHLH domain coding region, respectively. **b**, Sequence alignment of putative translational products of wild type (WT) and Mp*ice2*^*ge*^ mutants. Asterisks indicate the amino acids that are assumed to be important for binding to the E-box.

**Extended Data Fig. 9.**
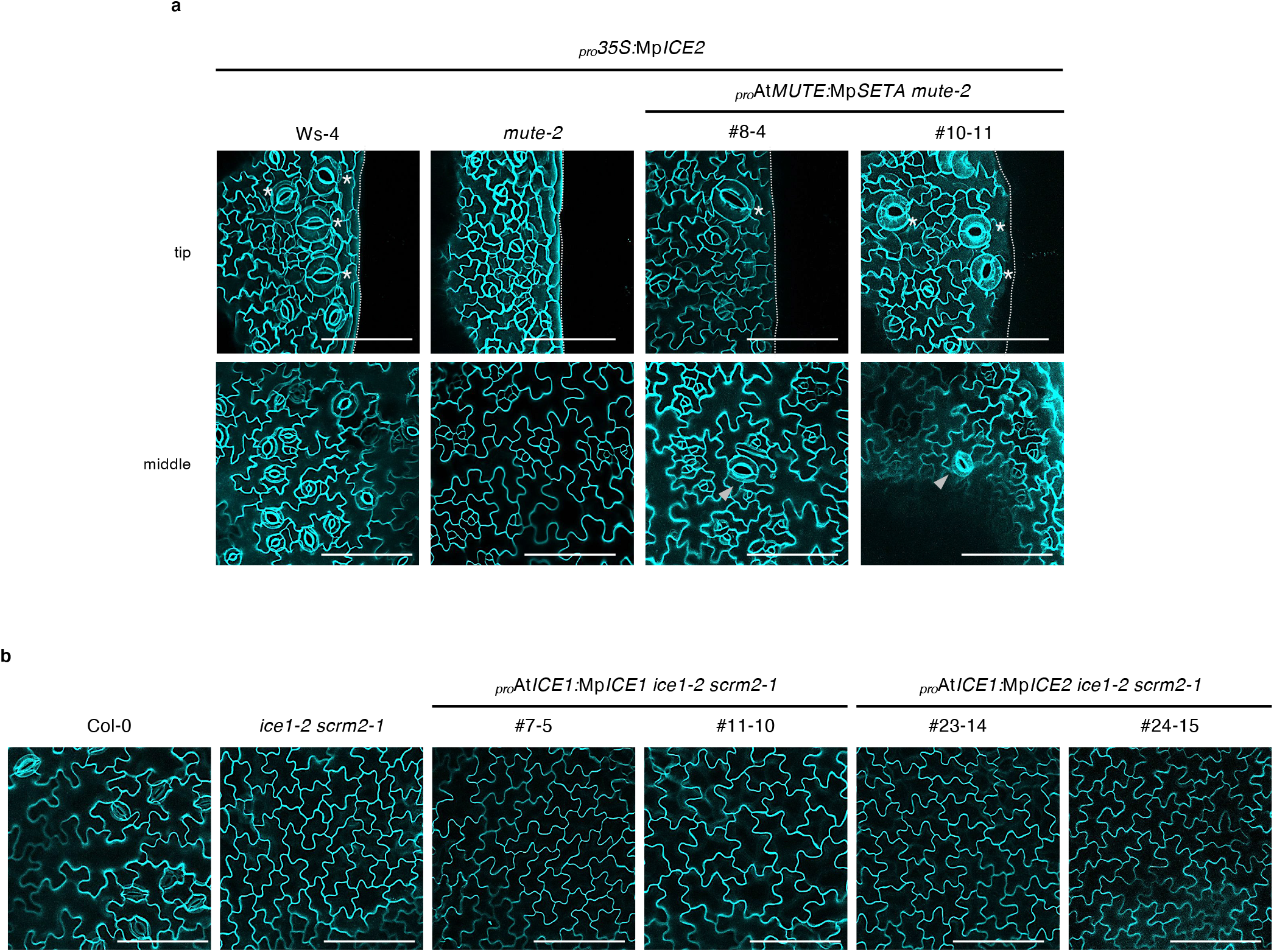
Functional analysis of Mp*ICE1* and Mp*ICE2* in *A. thaliana* mutants. **a**, Confocal images of *A. thaliana* abaxial cotyledons of wild type (Ws-4), *ice1-2 scrm2-1*, and _*pro*_At*MUTE:*Mp*SETA mute-2* expressing Mp*ICE2* at 9 DAS. Arrowheads and asterisks indicate stomata and hydathode pores, respectively. **b**, Confocal images of *A. thaliana* abaxial leaves of wild type (Col-0), *ice1-2 scrm2-1*, _*pro*_At*ICE1:*Mp*ICE1 ice1-2 scrm2-1*, and _*pro*_At*ICE1:*Mp*ICE2 ice1-2 scrm2-1* at 13 DAS. Bars, 100 μm.

## Supplementary Information

Supplementary Tables 1 and 2

