## Supplementary_Tables for "Stomatal regulators are co-opted for seta development in the astomatous liverwort *Marchantia polymorpha*"

**Supplementary Table 1 | Cloning primers used in this study.**

| **Plasmid** | **Primer Name** | **Sequence (5’>3’)** | **Source** |
| --- | --- | --- | --- |
| pENTR1A-Mp*SETA* CDS | MpSETA_CDS_ENTR_F | atccggtaccgaattcgcATGGACGGTGTTGTTCTGGACGATG | this study |
|  | MpSETA_CDS_ENTR_R | gtgcggccgcgaattctaACCCATGTTCTCAAATTGAAGATTG | this study |
| pENTR1A-Mp*ICE2* CDS | MpICE2_CDS_ENTR_F | atccggtaccgaattcgcATGGCTTCGACCAGAGT | this study |
|  | MpICE2_CDS_ENTR_R | gtgcggccgcgaattctaCTGAAGAATCACATTTTGCTCT | this study |
| pENTR1A-*_pro_*Mp*SETA* | MpSETApro_ENTR_F | aaccaattcagtcgacCAAGCCAGGTGAAAAAAATC | this study |
|  | MpSETApro_ENTR_R | gtgcggccgcgaattCCGAAGCGATTGGACCAGTAT | this study |
| pENTR1A-*_pro_*Mp*ICE2:Citrine-GUS-NLS* | MpICE2pro_ENTR_F | atccggtaccgaattcgcACATTCCTACTTGCAGCACTGA | this study |
|  | MpICE2pro_Cit_R | gcccttgctcaccatGAGAGAGTGAGCGCCTTCG | this study |
|  | Citrine_proMpICE2_F | GGCGCTCACTCTCTCatggtgagcaagggcgag | this study |
|  | Citrine_GUS_R | acgtaacatctgcagcttgtacagctcgtccatgc | this study |
|  | GUS_Citrine_F | gacgagctgtacaagctgcagatgttacgtcctgt | this study |
|  | GUS_NLS_ENTR_R | gtgcggccgcgaattctatcctccaacctttctcttct | this study |
| pENTR1A-*g*Mp*SETA* | MpSETApro_ENTR_F & MpSETA_CDS_ENTR_R | see above | this study |
| pENTR1A-*g*Mp*ICE2* | MpICE2pro_ENTR_F & MpICE2_CDS_ENTR_R | see above | this study |
| pDEST-GBKT-Amp^r^ | AmprPro_SfoI | TGTCAGCGCAGGGGCGCCCGCGGAACCCCTATTTGTTT | this study |
|  | AmprCDS_SfoI | CAAAAAGAACCGGGCGCCTTACCAATGCTTAATCAGTGAGGC | this study |
| pENTR1A-At*ICE1* | AtICE1_CDS_ENTR_F | aaccaattcagtcgacATGGGTCTTGACGGAAACAA | this study |
|  | AtICE1_CDS_ENTR_R | aagctgggtctagatatccTCAGATCATACCAGCATACCCTG | this study |
| pENTR1A-At*SCRM2* | AtSCRM2_CDS_ENTR_F | aaccaattcagtcgacATGAACAGCGACGGTGTTTG | this study |
|  | AtSCRM2_CDS_ENTR_R | aagctgggtctagatatccTCAAACCAAACCAGCGTAACC | this study |
| pENTR 5’/TOPO-*_pro_*At*MUTE* | AtMUTEpro_F | CAGCAATTTGAAAAATCCA | this study |
|  | AtMUTEpro_R | gatacttaattgatcaagata | this study |
| pENTR 5’/TOPO-*_pro_*At*FAMA* | AtFAMApro_F | cttccaaacttcatgtatgaa | this study |
|  | AtFAMApro_R | tgctattcgtggtagttgata | this study |
| pENTR1A-*_pro_*At*SPCH:*Mp*SETA* | SPCHpro_ENTR_F | aaccaattcagtcgacAGATCATCACTGCGATAAGG | this study |
|  | SPCHpro_Mp_R | AACAACACCGTCCATCGTGATTAGAGATATATCCT | this study |
|  | MpSETA_SPCHpro_F | ATCTCTAATCACGATGGACGGTGTTGTTCTGGACGATG | this study |
| pENTR1A-*_pro_*At*ICE1:*Mp*ICE1* | proICE1_ENTR_F_Sal1 | aaccaattcagtcgacGGACCACCGTCAATAACATCG | this study |
|  | proICE1_MpBHLH16_R | TCTGGTTGTAGAAGTCATCATGCCAAAGTTGACACCTTTACC | this study |
|  | MpBHLH16_proICE1_F | gtaaaggtgtcaactttggcATGATGACTTCTACAACCAGA | this study |
|  | MpBHLH16_CDS_ENTR_R | gtgcggccgcgaattctaCTGAAGGGCACTGACGTTA | this study |
| pENTR1A-*_pro_*At*ICE1:*Mp*ICE2* | proICE1_MpBHLH17_R | CTGACTCTGGTCGAAGCCATGCCAAAGTTGACACCTTTACC | this study |
|  | MpBHLH17_proICE1_F | gtaaaggtgtcaactttggcATGGCTTCGACCAGAGT | this study |
| pJHY-TMp1-MpSETAko | MpSETA-KO-Pac-F | ctaaggtagcgattaatACTCCGAATTTAAACGAT | this study |
|  | MpSETA-KO-Pac-R | gcccgggcaagcttaatCCGGTAGCTTTTATTCGC | this study |
|  | MpSETA-KO-Asc-F | taaactagtggcgcgCTGCAAGATCAAAAGAAT | this study |
|  | MpSETA-KO-Asc-R | ttatccctaggcgcgTGACAATGCTTCGATCAC | this study |

**Supplementary Table 2 | Genotyping, RT-PCR, and genome-editing primers used in this study.**

| **Purpose** | **Name** | **Sequence (5’>3’)** | **Source** |
| --- | --- | --- | --- |
| For genotyping sequencing (Mp*seta^ko^*) | MpSETA_KO_F | AACAAAAAAGAAGCGAATAAAAGCTACCGG | this study |
|  | MpSETA_KO_R | AGCGATGCTTGATTCTTTTGATCTTGCAG | this study |
|  | MpSETA_5’genomic_F | TGTTTTCATTCTCTCTATGACTTTTCAAGT | this study |
|  | MpSETA_3’genomic_R | TATCTGGGTTTTCTAGAGCTCATATAAACT | this study |
|  | P1R | GAAGGCTTCTGATTGAAGTTTCCTTTTCTG | Ishizaki et al.^1^ |
|  | H1F | GTATAATGTATGCTATACGAAGTTATGTTT | Ishizaki et al.^1^ |
| For sex diagnosis | rbm27-F | ACTTTTGCAACAGCGACTTC | Fujisawa et al.^2^ |
|  | rbm27-R | GCCTGCAATATAGCCTTCAA | Fujisawa et al.^2^ |
|  | rhf73-F | GAACCCGAAACTCAGGTTTT | Fujisawa et al.^2^ |
|  | rhf73-R | ATAACAGCCAAACGGATCAA | Fujisawa et al.^2^ |
| For RT-PCR | MpSETA_RT_F | ATGGACGGTGTTGTTCTGGACGATG | this study |
|  | MpSETA_RT_R | ACCCATGTTCTCAAATTGAAGATTG | this study |
|  | MpEF1α_F | AGGTTGTCACCATGGGAAAGGAGA | Rövekamp et al.^3^ |
|  | MpEF1α_R | TCACACGCTTGTCAATACCTCCCA | Rövekamp et al.^3^ |
| For genome editing (Mp*ice2^ge^*) | MpICE2_sgRNA1_F | CTCG CTTGCCTGCGAAGAATCTCA | this study |
|  | MpICE2_sgRNA1_R | AAAC TGAGATTCTTCGCAGGCAAG | this study |
|  | MpICE2_sgRNA2_F | CTCG GGACAGAGCATCAATCCTTG | this study |
|  | MpICE2_sgRNA2_R | AAAC CAAGGATTGATGCTCTGTCC | this study |

1.	Ishizaki, K., Johzuka-Hisatomi, Y., Ishida, S., Iida, S. & Kohchi, T. Homologous recombination-mediated gene targeting in the liverwort *Marchantia polymorpha* L. *Sci. Rep.* **3**, 1–6 (2013).

2.	Fujisawa, M. *et al.* Isolation of X and Y chromosome-specific DNA markers from a liverwort, *Marchantia polymorpha,* by representational difference analysis. *Genetics* **159**, 981–985 (2001).

3.	Rövekamp, M., Bowman, J. L. & Grossniklaus, U. *Marchantia* Mp*RKD* Regulates the Gametophyte-Sporophyte Transition by Keeping Egg Cells Quiescent in the Absence of Fertilization. *Curr. Biol.* **26**, 1782–1789 (2016).
